# Effects of land-use change and related pressures on alien and native subsets of island communities

**DOI:** 10.1101/2019.12.16.878041

**Authors:** Katia Sánchez-Ortiz, Kara J. M. Taylor, Adriana De Palma, Franz Essl, Wayne Dawson, Holger Kreft, Jan Pergl, Petr Pyšek, Mark van Kleunen, Patrick Weigelt, Andy Purvis

## Abstract

Island species and habitats are particularly vulnerable to human disturbances, and anthropogenic changes are increasingly overwriting natural island biogeographic patterns. However, quantitative comparisons of how native and alien assemblages respond to human disturbances are scarce. Using data from 6,242 species of vertebrates, invertebrates and plants, from 7,718 sites on 81 islands, we model how land-use change, human population density and distance to the nearest road affect local assemblages of alien and native species on islands. We found that land-use change reduces both richness and abundance of native species, whereas the number and abundance of alien species are high in plantation forests and agricultural or urban sites. In contrast to the long-established pattern for native species (i.e., decline in species number with island isolation), more isolated islands have more alien species across most land uses than do less isolated islands. We show that alien species play a major role in the turnover of island assemblages: our models show that aliens outnumber natives among the species present at disturbed sites but absent from minimally-disturbed primary vegetation. Finally, we found a homogenization pattern for both native and alien assemblages across sites within most land uses. The declines of native species on islands in the face of human pressures, and the particular proneness to invasions of the more remote islands, highlight the need to reduce the intensity of human pressures on islands and to prevent the introduction and establishment of alien species.

## Introduction

Land use and related anthropogenic pressures – such as habitat destruction, use intensity and human population growth – strongly modify species diversity within local ecological assemblages [1–3] and turnover between them [4], in ways that can differ significantly between ecological systems or geographic regions [1,5,6]. Islands – landmasses completely surrounded by ocean and smaller than continents – represent a set of ecosystems and regions where the effects of these pressures may be particularly severe, for two reasons. First, many species native to islands have characteristics – e.g., narrow-range endemism, small population size – that make them both irreplaceable and particularly vulnerable to human disturbances [7–9]. Second, islands have been highlighted as hotspots of alien species [10–13] and as systems at higher risk of invasions [14–17].

Natural and anthropogenic factors may combine to make islands, and their site-level communities, prone to invasion. Islands – especially remote ones – have a relatively small pool of native species [18] from which their communities are assembled. The resulting low species richness for their area [19] is in turn hypothesised to result in more available resources [15], low pressures from predators or pathogens, and disharmonic communities (i.e. with a biased representation of certain taxonomic or functional groups) [16,20]. Additionally, many native species have low competitive ability [16] and have evolved reduced dispersal abilities [20] and reproductive output [21–23]. These natural factors combine to facilitate establishment by alien colonists, especially in disturbed sites [24–26]. Land-use change involves site-level disturbance, favouring the establishment of alien species (often good dispersers that reproduce rapidly and tolerate a broad range of conditions [27–29]). Additionally, humans have directly introduced many alien species to islands, with propagule pressure (i.e., number of released individuals [30]) and the presence of other groups of introduced species [31] both influencing the probability of establishment.

Anthropogenic disturbances favour alien species, but disturbed habitats can represent novel environments for some native species, to which they may not be sufficiently well adapted to exploit [15]. Natives with traits related to a high extinction risk – such as large size [32,33], low fecundity, limited dispersal abilities, high habitat specificity [34] and small range-size [33,35] – are particularly unlikely to tolerate human impacts, whereas those with the opposite traits are likely to be more resilient. Both the addition of aliens and the accelerated turnover of natives can drive changes in among-site species turnover, including biotic homogenization [36]: their relative importance remains an open question. Quantitative comparisons of how native and alien species respond to human disturbances on islands are still scarce (but see [24,25,37,38]).

If alien species follow natural island biogeographic patterns, large islands and islands close to other land masses would have more alien species than small and isolated ones [39]. However, evidence is mounting that anthropogenic processes have already altered fundamental biogeographical patterns such as the species–isolation relationship [12,40–42]. Anthropogenic factors such as colonisation pressure (i.e., number of species introduced to a defined location [11,43]) or economic isolation of islands [41] may have become more important for the assembly and diversity of insular biota than geographic isolation. Measures of economic activity are related to colonisation [44] and propagule pressures (e.g., trade volumes), ecological disturbances [45] and infrastructure development [46] (e.g., roads), all of which ease the arrival and establishment of alien species [45,46]. Here, as a simple proxy for islands’ economic connectance, we use per-capita Gross Domestic Product (GDP), which has previously been shown to correlate across countries globally with the numbers of alien introductions [11] and, among countries within regions, with numbers of established alien plants [47], amphibian and reptile species [10].

In this paper, we use data from 6,242 species (including vertebrates, invertebrates and plants), from 7,718 sites on 81 islands in the PREDICTS database [48], whose native/alien status could be determined, to model how land-use change and related pressures affect local native and alien assemblages on islands. This study is the first global analysis of its kind to include a wide range of taxa while focusing specifically on islands. Based on models of site-level total abundance and species richness, we show that three human pressures (land use, human population density and distance to the nearest road) affect alien assemblages differently from native assemblages. Additionally, we test whether richness and abundance of alien species is predicted by measures of island size, geographic isolation and economic connectance. To evaluate the turnover of native and alien assemblages caused by land-use change on islands, we estimate how species composition of alien and native communities in minimally-disturbed sites is affected by land-use change. Finally, for native and alien assemblages separately, we estimate the homogeneity of assemblages among sites within the same land use.

## Methods

### Biodiversity data and species status

All data on species abundance and occurrence at sites on islands (including Australia) were extracted from the PREDICTS database [48] in October 2016. The database collates data from published research that compared local biodiversity across sites in different land uses, and is structured hierarchically into Data Sources (publications), Studies (different sampling methods within a source), Blocks (spatial blocks, if present in the study) and Sites [49]. Australia was treated as an island because of its island-like characteristics – e.g., long isolation history (complete isolation from other continents ∼ 33 Mya [50]) and markedly smaller size than other continents. The name of the island where each site was located was determined by matching the site coordinates with the Global Island Database, ver. 2.1 [51]. All data processing and statistical analyses were performed using R Version 3.3.3 [52], unless otherwise stated.

The island data from PREDICTS included 17,776 species and 1,339,339 biodiversity records (each being a single abundance measure or occurrence record of a species at a site within a study). Only 42 of the168 data sources had already classified the sampled species as aliens or natives at the sites sampled. To classify as many remaining species as possible, we matched PREDICTS records with 27 external sources of information on native and alien ranges (Table 1 in S1 File). Three global databases provide island-specific (rather than country-level) species classifications: the Global Naturalized Alien Flora database (GloNAF; [53,54]), the Global Inventory of Floras and Traits (GIFT; [55]) and the Threatened Island Biodiversity database (TIB; [56]). We identified all matching species-island combinations between these three databases and the PREDICTS island data. For remaining taxa, we queried additional databases (Table 1 in S1 File) as well as GloNAF for species status at a country level, via either downloadable datasets or an Application Programming Interface (API). Most of these external sources provided a direct indication of the species status in different locations (e.g. alien, invasive, naturalized, native, endemic). When databases provided geographic ranges for species, we first extracted the status for the matching species-country combinations, but also classified species according to the geographic ranges provided by these sources: i.e., species whose sampling location in our dataset was not included in their native range were classified as alien. In a final step, we searched for data sources that could help classify species with many records in our data set that had remained unclassified to this point (mainly arthropods); these additional sources included taxonomic experts from different institutions, publications and databases for specific taxa and countries.

### Statistical modelling

#### Models of total abundance and species richness

For each site, we split the PREDICTS data into native species and alien species, discarding data that could not be classified. For aliens and natives separately, we calculated each site’s total abundance (sum of abundances of all taxa present) and species richness (number of unique taxa present). When sampling effort (provided in source publications) varied among sites within a study, total abundance was divided by the sampling effort to make data comparable among the study’s sites; this was the case for 24 of 163 studies. Not all the studies reported abundance data, meaning that models of abundance were based on 18 fewer studies. Dropping these 18 studies from the richness dataset (to use the same data in both models) did not markedly change our results; therefore, we present the results from the model using all available data for species richness.

There were 74 studies that reported data only for natives (66 studies) or aliens (8 studies). Referring back to the published methodologies for each such study, thirteen studies had explicitly targeted only native species and two only aliens; most of these only focused on sampling a few (between one and eight) species instead of native or alien communities *per se*. For sites within these studies, we treated the site-level diversity of the unsampled class as missing. The remaining studies had targeted the entire assemblage but the recorded species whose status could be determined were either all natives (53 studies) or all aliens (6 studies). For sites within these studies, we assigned zero site-level diversity metrics (for metrics originally available in the study) for the missing group in the study (aliens or natives), To account for the fact that not all species in the studies could be classified as aliens or natives, we weighted sites in the analyses that follow, with each site’s weight being its proportion of classified species (i.e., adding weight to sites which kept data for most of their originally sampled species).

The studies in the PREDICTS database differ greatly in their sampling methods, taxonomic focus and locations (Tables 4 and 5 and Fig 1 in S1 File); and some studies have spatial blocks or split-plot designs. To accommodate the resulting heterogeneity and non-independence, we fitted mixed-effect models (using the ‘*lme4’* package ver. 1.1-15 – [57]) to model the responses of alien and native species richness and total abundance to human pressures. As fixed effects, the full models included two three-way interactions between two human pressures at a site-level and the species status (alien/native): one between land use, human population density (HPD) and species status; and the second between land use, distance to nearest road (DistRd) and the species status. Because not all abundance data were counts of individuals (e.g., relative abundance), Poisson or quasipoisson models could not be used: instead, sites’ total abundance was rescaled to a zero-to-one scale within each study (dividing by the maximum abundance within the study) to reduce the variance among studies, then square-root transformed (which gave a better residual distribution than ln(x+1) transformation), and modelled using a Gaussian error structure. Species richness (always a count) was modelled using a Poisson error structure and log-link function.

**Fig 1.**
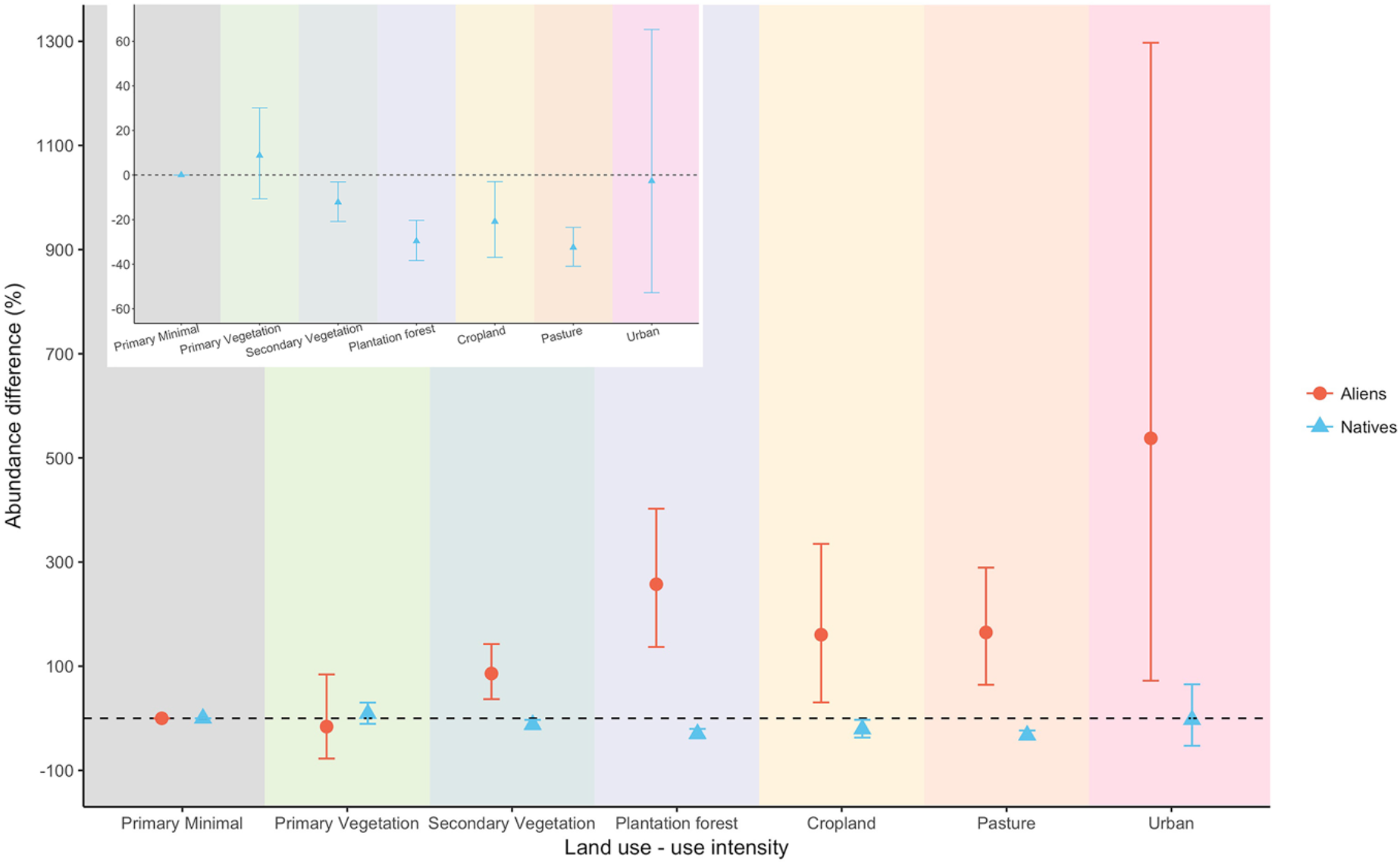
Response of total abundance of aliens and natives to land use. Values indicate decrease or increase in percentage of total abundance using minimally-used primary vegetation as baseline (dashed line). Bars indicate 95% confidence intervals, which are asymmetric because of back-transformation from the square-root scale. The inset expands the y-axis for natives, which are not clearly visible in the main plot.

Site-level data for human population density (for the year 2000) were taken from the Global Rural-Urban Mapping Project, ver. 1 (GRUMPv1) [58] at a resolution of 30 arc sec (∼1 km). Site distance to the nearest road came from the Global Roads Open Access Data Set, ver. 1 (gROADSv1) [59]; we projected this vector map of the world’s roads onto an equal-area (Behrmann) projection and calculated the average distance to the nearest road within each 782-m grid cell using the ‘*Euclidean Distance*’ function, from which we produced a gridded map of distance to nearest road (using Python code written for the *arcpy* module of ArcMap ver. 10.3), which we reprojected back to a WGS 1984 projection at 30 arc sec resolution. HPD and DistRd for each site was obtained by matching the sites’ coordinates with these gridded maps. We treated HPD and DistRd as constant within grid cells. HPD and DistRd were ln transformed (ln(x+1) in the case of HPD) to reduce skew and rescaled to a zero-to-one scale (to reduce collinearity) prior to modelling, and fitted as quadratic orthogonal polynomials.

Sites in the PREDICTS database had previously been classified into 10 land-use categories and three land-use intensities (minimal, light and intense) within each land use [49]. For our models, these categories were collapsed into seven final land-use/use-intensity classes: 1) Minimally-used primary vegetation (henceforth PriMin; used as a baseline representing minimally disturbed sites), 2) Primary vegetation (other than minimal use), 3) Secondary vegetation, 4) Plantation forest, 5) Cropland, 6) Pasture and 7) Urban. These collapsed classes gave reasonable sample sizes (at least 300 sites) within each land-use class for both alien and native data (Table 7 in S1 File). Sites missing data for any of the human pressures were excluded from these models.

As random effects for the models, we initially considered random-intercept terms of identity of the study from which data came from (to account for the variability in sampled biodiversity, sampling methods and location across studies), identity of the spatial block — nested within study — in which the site was located (to account for the spatial arrangement of sites) and island (to account for the variability of sampled biodiversity across islands). We also considered random slopes to account for the variation among studies in the relationships between sampled biodiversity and land use/use intensity. To identify the best random-effects structure, we compared four possibilities following the procedure in [60]: (i) random slopes for land uses within study + random intercepts of study, block nested within study and island, (ii) same structure as (i) but dropping the island random intercept, (iii) random intercepts of study, block nested within study and island, and (iv) random intercepts of study and block nested within study. For the richness model, we added an observation-level random effect (i.e. site identity), to account for overdispersion [61]. Once the random-effects structure had been chosen, the fixed-effects structure of the final models was determined using backwards stepwise model simplification and likelihood ratio tests with models fitted using maximum likelihood [60]. Generalized Variance Inflation Factors (GVIFs) [60] did not indicate strong collinearity among the explanatory variables and model diagnostics showed that the final abundance and richness models fulfilled homogeneity and normality assumptions (Fig 2 in S1 File).

**Fig 2.**
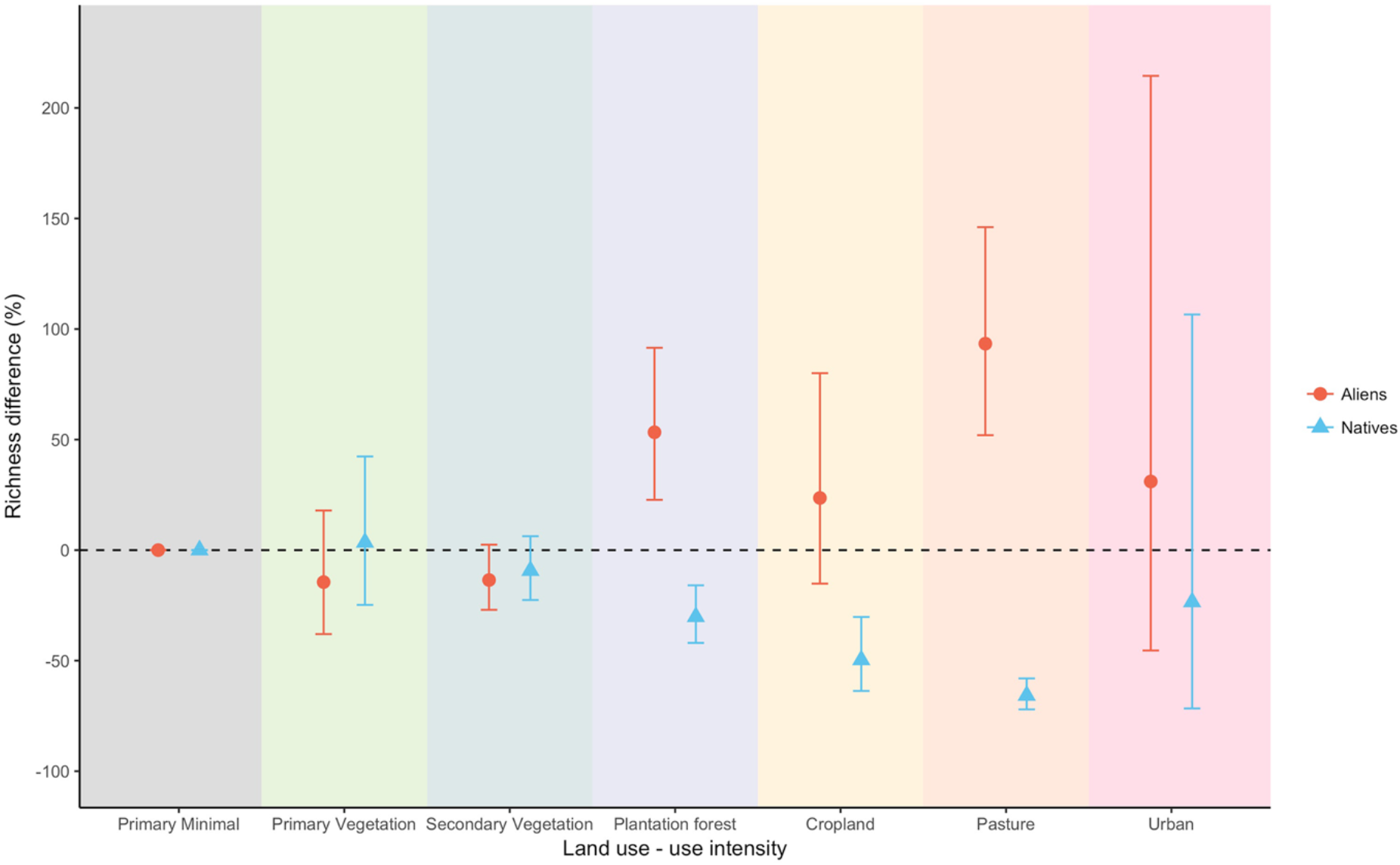
Response of richness of aliens and natives to land use. Values indicate decrease or increase in percentage of species richness using minimally-used primary vegetation as baseline (dashed line). Bars indicate 95% confidence intervals and are asymmetric because of back-transformation from the logarithmic scale of the models.

#### Models including island characteristics as predictors

We tested whether any of three island characteristics – area, isolation and GDP per capita – predicted total abundance and richness of aliens. Island area (in km^2^) was calculated using a global layer of land polygons taken from OpenStreetMap [62] in cylindrical equal area projection. As a proxy for island isolation, we used Weigelt et al.’s [63] values for the sum of the proportions of landmass within buffer distances of 100, 1,000 and 10,000 km around the island (henceforth, surrounding landmass), which has been shown to predict diversity of plants on islands at a global scale [64]. Data for three Japanese islands and Australia (Table 11 in S1 File) were missing from Weigelt et al. [63], so these four islands were not included in the surrounding landmass models. GDP per capita (in current US dollars for the year 2005) for each site’s country came from World Bank Open Data.

Using only the alien data, we fitted six mixed-effects models – testing each of the three island characteristics in turn as predictors of richness and abundance – with the same modelling approach as used above with the following exceptions. We dropped studies that had zero site-level diversity metrics for aliens across all sites, which resulted from adding these metrics to some studies missing alien data. As fixed effects, each initial model included one two-way interaction between land use and the island characteristic. To reduce skewness and improve the distribution of residuals [65], island area and GDP per capita were ln-transformed. Surrounding landmass did not require transformation. We considered two possible random-effects structures for the models: (i) random intercepts of study (ii) random intercepts of study and island. The richness models were not overdispersed. Based on GVIFs, we did not find strong collinearity among the explanatory variables of any of the models. We performed post hoc analysis (package ‘*phia*’ ver. 0.2-1 – [66]) to test whether the coefficients of the interaction between the land uses and the island trait were significantly non-zero.

#### Models of compositional similarity

To model how land-use change affects the turnover of native and alien assemblages on islands, we model pairwise differences in assemblage composition among sites using a modified matrix regression, explained in more detail below. Under this approach, the non-independence among the pairwise comparisons necessitates the use of permutation tests to assess statistical significance [67].

We first calculated compositional similarity (for aliens and natives separately) for all possible pairwise comparisons between sites (including both forward and reverse comparisons) within each study using the abundance-based (*J*_*A*_) and richness-based (*J*_*R*_) asymmetric Jaccard Index, calculated as: *J*_*R*_= *S*_*ij*_ /*S*_*j*_, where *S*_*ij*_ is the number of species common to both sites and *S*_*j*_ is the number of species in site *j*; *J*_*A*_= *A*_*ij*_ /*A*_*j*_, where *A*_*ij*_ is the summed abundance at site *j* of all species common to both sites and *A*_*j*_ is the summed abundance of all species at site *j* [68]. These asymmetric measures reflect the possibility of one site’s assemblage being largely nested within another [69]. Both measures are one if all taxa at site *j* are also present at site *i*, zero if the two sites share no taxa in common, and undefined (and dropped from analyses) if neither site had any organisms sampled. Low *J*_*A*_ and *J*_*R*_ values mainly result from a high number or high abundance of novel or unique species in site *j* (i.e., species recorded at site *j* but missing in site *i*).

We fitted mixed-effects models to model both *J*_*A*_ and *J*_*R*_ as a function of the sites’ land uses (henceforth land-use contrast; e.g, PriMin-Pasture), the geographic and environmental distance between the sites (to account for similarity decay with geographic [70] and environmental distance) and the assemblage status (native or alien). Land-use contrast was a 49-level factor (i.e., all possible combinations for the seven land-use categories); however, we only focus on results for 13 land-use contrasts (Table 16 in S1 File): – the set of contrasts with site *i* in PriMin, estimating how the composition of native and alien species of minimally-disturbed sites is affected by change to each other land use; and the set of contrasts with both sites in the same land use, estimating how similar communities are among sites facing similar pressures. For both alien and native data, the land-use contrasts of interest had sample sizes >1900 and included data from five or more studies (Table 16 in S1 File); the only exception was PriMin-Urban, for which data only came from one study for aliens and two studies for natives. Because estimates based on so few studies are unreliable [71], we do not report the results for this contrast (model coefficients for all land-use contrasts are presented in Table 17 in S1 File). Geographic distance between sites was calculated from the sites’ coordinates using the ‘*distHaversine*’ function in the ‘*geosphere*’ package ver. 1.5-7 [72]. Environmental distance was estimated as the dissimilarity between sites, measured as the Gower’s distance [73] between them calculated from site-level data for altitude and four bioclimatic variables (maximum and minimum temperature and precipitation of wettest and driest month) at 1-km spatial resolution [74]. We excluded pairwise comparisons for which environmental distance could not be calculated due to missing data or where geographic distance was zero as a result of coordinate imprecision. We also excluded studies that sampled a single species, where sampling effort varied among sites and that did not report abundance data so that our abundance- and richness-based models would use data from the same studies.

As fixed effects, the models included two-way interactions between assemblage status (alien/native) and each of geographic distance, environmental distance and land-use contrast. Study identity was included as a random intercept. The models were fitted adding weight to pairwise comparisons with a higher proportion of classified species in site *j*. We used the PriMin-PriMin contrast as the intercept level in the models and a baseline against which to compare the other land-use contrasts, since it represents the natural spatial turnover of species. Prior to modelling, compositional similarity was logit-transformed (using an adjustment of 0.01 to deal with 0 and 1 values [75]). To deal with extreme values [65] and normalise the data distribution, environmental distance was transformed using cube root, and geographic distance was first divided by the median maximum linear extent (maximum distance between sampling points within a site; e.g., extent of a single quadrat or the greatest distance between any two traps within a site) of the sites in the dataset (which reflects site size [76]) and then ln-transformed; as a result, a transformed geographic distance of zero corresponds to adjacent sites. GVIFs did not indicate strong collinearity among the explanatory variables of the models for *J*_*A*_ and *J*_*R*_, which also fulfilled homogeneity and normality assumptions (Fig 13 in S1 File).

Because these models use all pairwise comparisons, they have extensive pseudo-replication, precluding the use of standard statistical approaches to simplify the fixed effects of the models. We therefore used permutation tests [67] to assess significance of terms during backwards stepwise model simplification based on likelihood ratios. The model dataset was permuted 199 times by randomly shuffling the response variable data among sites within studies while holding data for all explanatory variables constant. The likelihood ratio between models at successive stages of model simplification (i.e., between a more complex and a less complex model) was compared against a null distribution of ratios of initial and reduced models fitted with the 199 permuted datasets (function ‘*as*.*randtest*’ in the ‘*ade4*’ package ver. 1.7-10 – [77]). The statistical significance of the model coefficients for interactions between alien/native status and the other explanatory variables was tested in the same way, except that the comparisons between the model coefficients and the coefficients obtained from the permutation trials were two-tailed.

## Results

3,059 species recorded from 2,840 island sites in the PREDICTS database were already classified as native or aliens by 42 data sources. There were 13,108 unique combinations of island and species with a Latin binomial, of which 1,109 (for 1,039 species) could be classified using the island-specific databases, and 10,968 unique combinations of species and country, of which 2,160 (for 1,866 species) could be classified using the country-level external sources. The resulting data included 794 alien species and 5,517 native species (Table 6 in S1 File). In total, we were able to classify ∼52% of the island biodiversity records in the PREDICTS database as native or alien (Table 4 in S1 File), and retained ∼70% of the island studies and ∼75% of island sites in the PREDICTS database. The final site-level diversity dataset included data for alien species from 7,474 sites in 150 studies, 54 islands and 27 countries, and data for native species from 7,524 sites, in 161 studies, 81 islands and 28 countries (location of sites is shown in Fig 1 in S1 File). The datasets used in modelling are described further in Tables 7, 11 and 16 in the S1 File and are available at https://doi.org/10.5519/0047472.

### Effects of human pressures on native and alien species

The best random-effects structure for the abundance and richness models included random slopes for land uses within study and random intercepts of study, block nested within study and island (Table 8 in S1 File); however, failure of richness models with this structure to converge forced us to model richness without the island random intercept. We chose to also model abundance without this random intercept to keep the same random-effects structure in both models (after confirming that results from abundance models with and without the island random intercept differed minimally). Aliens and natives responded very differently to human pressures, in terms of both overall abundance (Table 9 in S1 File) and species richness (Table 10 in S1 File).

On average, ∼17% of the species and individuals in sites in minimally-disturbed primary vegetation (in the species-richness and abundance datasets, respectively) were aliens (and ∼83% natives) (Figs 7 and 8 in S1 File). Compared with PriMin, four land uses had significantly lower overall abundance of natives: pastures (−32%), plantation forests (−30%), croplands (− 21%) and secondary vegetation (−12%) (Fig 1). By contrast, aliens were very much more abundant in all other land uses, especially in the human-dominated land uses, than in primary vegetation (Fig 1). Plantation forests and agricultural land uses (croplands and pastures) had markedly lower native species-richness, but much higher alien species-richness (plantation forests and pastures for the latter case), than seen in PriMin sites (Fig 2).

The covariates in these models – HPD and DistRd – interact significantly with land use to shape both abundance (Table 9 in S1 File) and species richness (Table 10 in S1 File) in ways that differ between natives and aliens, but effects are not even qualitatively consistent among land uses or between aliens and natives (Figure 3 to 6 in S1 File).

**Fig 3.**
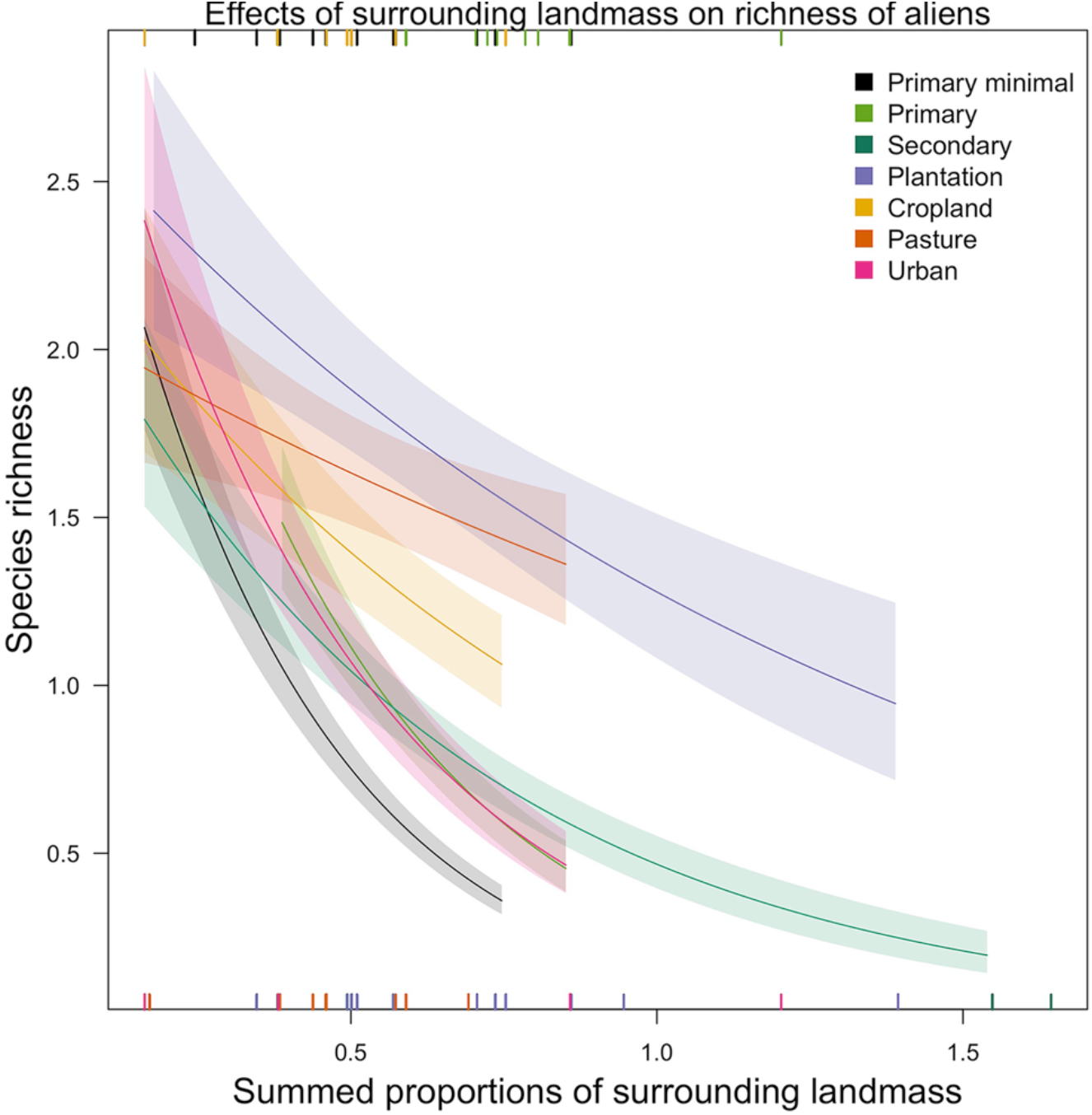
Effects of surrounding landmass on richness of alien species across land uses. For clarity, the error bars show half the standard error. The lines are curved because the response variable has been back-transformed from the logarithmic scale of the model. Slopes – i.e., estimates (est) – that are significantly different from zero: PriMin (est= −2.9, SE= 0.7, Chisq= 20.6, P= <0.001), Primary vegetation (est= −2.5, SE= 0.8, Chisq= 7.0, P= <0.01), Secondary vegetation (est= −1.6, SE= 0.4, Chisq= 6.9, P= <0.01) and Urban (est= −2.4, SE= 0.5, Chisq= 10.3, P= <0.01). Rugs along the horizontal margins show the values of surrounding landmass represented (across land uses) in the model data set (rugs for minimally-used primary vegetation, primary vegetation and croplands in the top margin and the rest of the land uses in the bottom margin). Rugs for land uses overlap, so some data are not visible.

### Island characteristics as predictors of alien abundance and richness

For all six models relating alien abundance or species richness to island characteristics, the best random-effects structure included random intercepts of study and island (Tables 12 and 14 in S1 File). None of the models could be simplified – land use interacted significantly with each island characteristic (Tables 13 and 15 in S1 File). However, only a few land uses had significant interaction coefficients (Figs 10 and 12 in S1 File), especially in the richness models, and hence few clear patterns could be discerned. The clearest pattern was seen in the effects of surrounding land mass on site-level species richness (Fig 3): sites within most land uses (but not croplands, pastures or plantation forests) have more alien species on isolated islands (i.e., those with the least surrounding landmass) than on less isolated ones. This trend was particularly strong in PriMin, primary vegetation and urban sites.

### Land use and compositional similarity of native vs alien assemblages

All interactions were significant in the abundance-based model (P= 0.005 for all interaction terms, meaning that none of the 199 permutation trials produced any interaction as strong as those in the actual data). In the richness-based model, only environmental distance did not interact significantly with the alien/native term (P= 0.24); the minimum adequate model retained all other interactions and environmental distance as a main effect (P= 0.005 in all cases). For both *J*_*A*_ and *J*_*R*_, the decline in similarity with geographic distance (i.e., distance-decay) is significantly steeper for alien assemblages than for native ones; for *J*_*A*_, the reverse is true for the decline with environmental distance (Table 17 and Fig 14 in S1 File).

Once distance-decay effects are controlled for, land use affects compositional similarity to PriMin (i.e., the presence and abundance of novel species in land uses other than PriMin) significantly more in alien than native assemblages, for both *J*_*R*_ and *J*_*A*_ (Fig 4; Fig 15 in S1 File). Alien assemblages of PriMin sites were more compositionally similar to each other than to those in other land uses, especially agriculture. In contrast, native assemblages of PriMin sites were slightly less similar to each other than to assemblages of sites in most other land uses.

**Fig 4.**
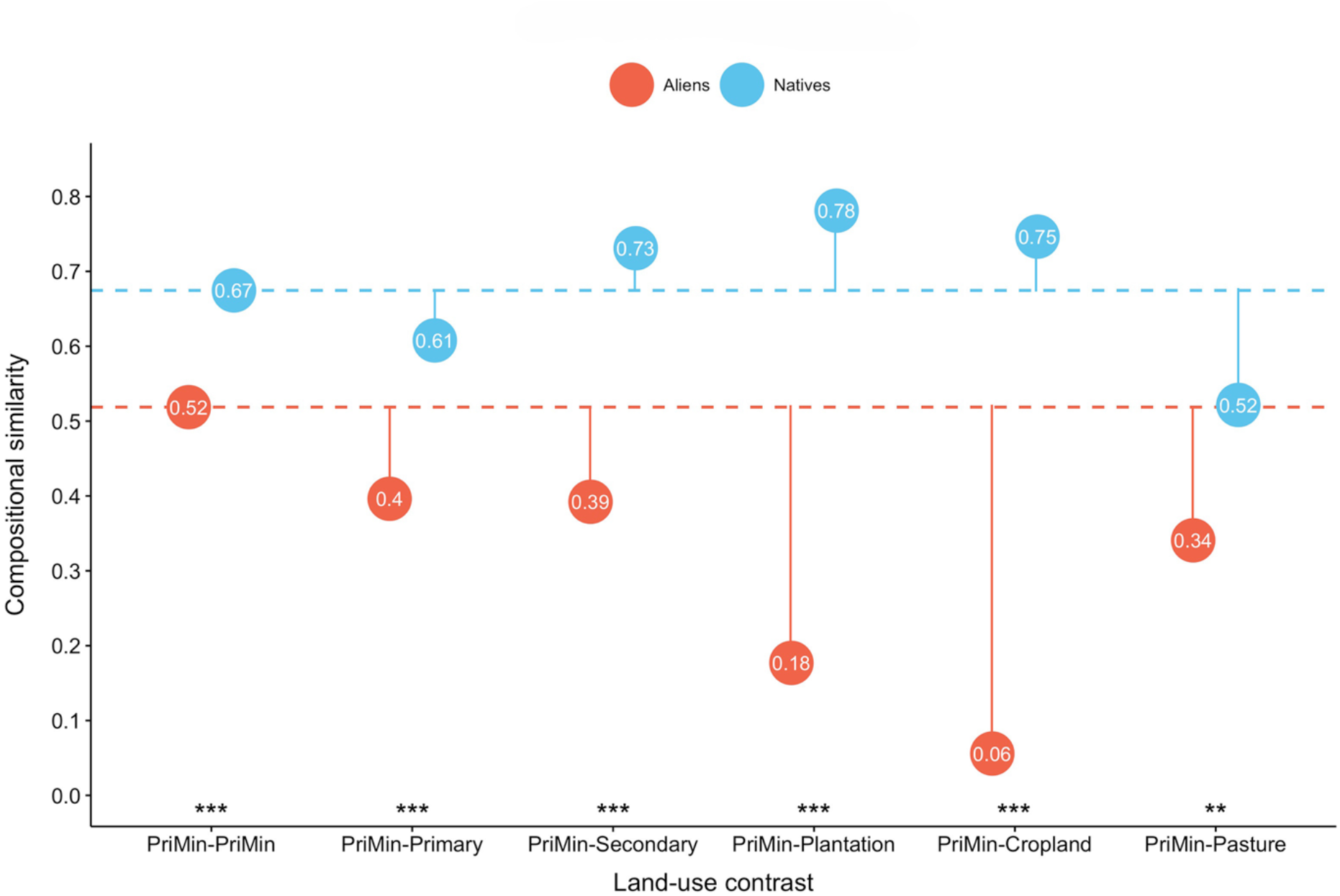
Richness-based (*J*_*R*_) compositional similarity estimates for land-use contrasts where site *i* is in PriMin. Solid lines show the magnitude of change in *J*_*R*_ driven by change to different land uses; the baseline is compositional similarity between PriMin sites for alien and native assemblages respectively (dashed lines). Significance (indicated by stars) is shown for alien/native differences for *J*_*R*_ changes from PriMin-PriMin on a logit scale (results from permutation tests and two-tailed tests comparing the coefficients for interaction between alien/native and land-use contrast to null distributions). Results for the PriMin-Urban contrast are not shown because sample sizes for this contrast were very small (for these coefficients, see Table 16 in S1 File) Significance codes: <0.05**, 0.005***

Moving to compositional similarity within each land use, pairs of sites within most land uses tend to have more similar assemblages than do pairs of PriMin sites, for both alien and native assemblages and for both *J*_*R*_ and *J*_*A*_ (Fig 5; Fig 16 in S1 File). Most of these within-land-use similarities differed significantly between alien and native assemblages; the reduction of spatial beta diversity (when compared to similarity between PriMin sites) is stronger for alien assemblages in plantation forests, but also, for *J*_*R*_, in urban sites. Reduction of spatial beta diversity was stronger in native than alien assemblages in models for *J*_*R*_ in secondary vegetation and cropland but only slightly so (Fig 5), and in models for *J*_*A*_ in primary and secondary vegetation and pastures (Fig 16 in S1 File).

**Fig 5.**
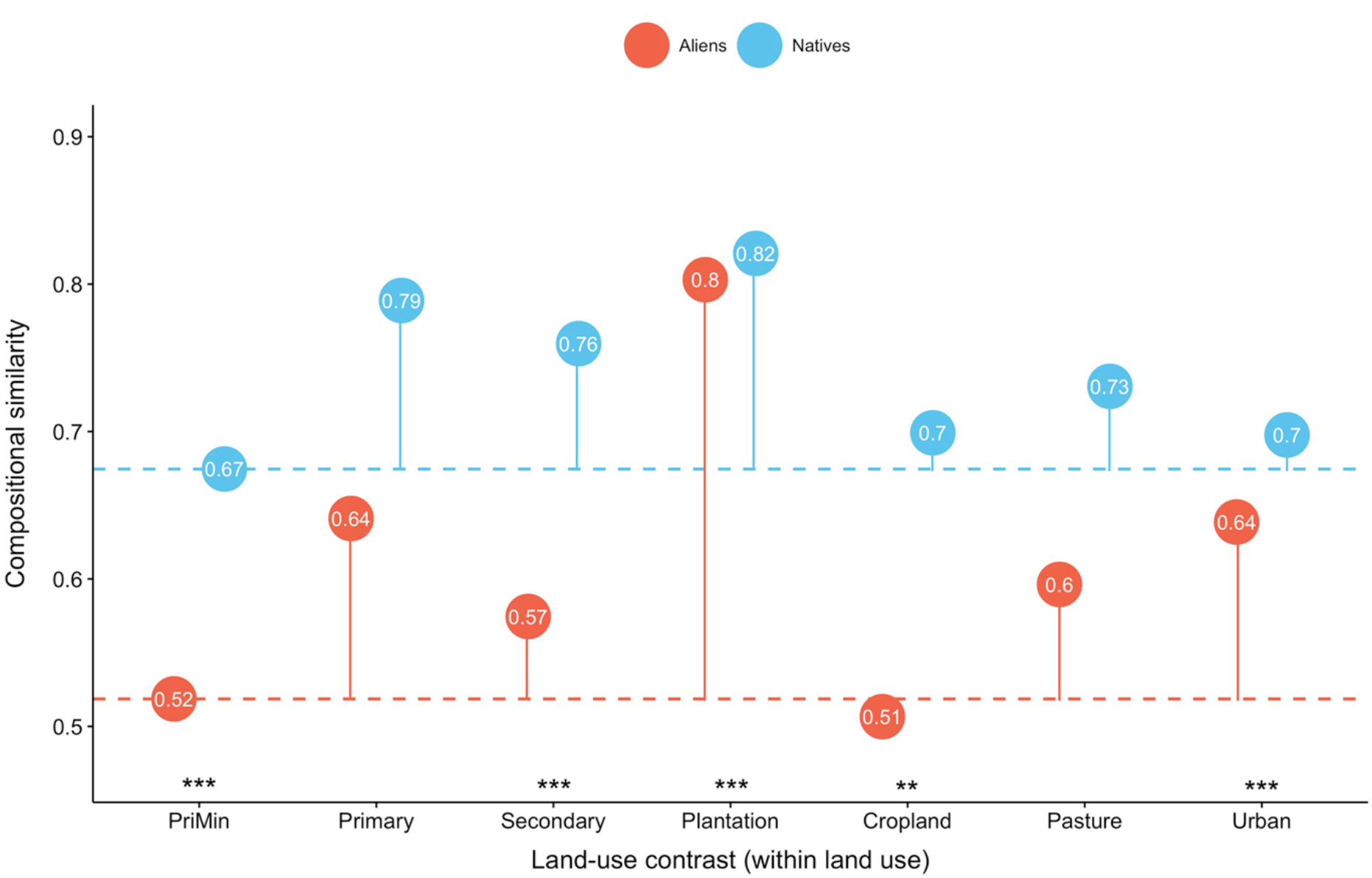
Richness-based (*J*_*R*_) compositional similarity estimates for alien and native assemblages in sites within the same land use. Each category corresponds to a land-use contrast (i.e., Cropland= Cropland-Cropland). Solid lines show the magnitude of change in *J*_*R*_ using PriMin-PriMin compositional similarity as baseline (dashed lines). Significance connotation and codes as in Fig 4.

## Discussion

Human-dominated land uses markedly decrease the abundance and richness of native species in island sites and at the same time dramatically increase the abundance and richness of alien species. These findings are in line with previous suggestions that habitat modification tends to have particularly negative impacts on native [34,78] and narrow-range species [79], and with the hypothesis that alien species can be very successful in disturbed habitats [15,80– 82]. Our results also corroborate studies that identified agriculture as a main cause of biodiversity loss [83,84]: cropland and pasture sites showed the largest reductions in native species-richness (with drastic reductions of −50% and −65% respectively) and, in pastures, abundance (−30% reduction).

On average, ∼17% of the species and total abundance in minimally-disturbed sites corresponded to alien species. The presence of so many aliens even in sites with the least human impact violates the assumption that such sites accurately represent the assemblages of disturbed sites prior to land-use change – the central assumption of space-for-time substitution – with the result that statistical models such as ours are likely to underestimate the net effects of land-use change on local assemblages on islands [85]. Whether this problem extends to mainlands depends on the proportion of aliens in minimally-impacted mainland sites, which we are not able to estimate from our data. The proportion might be either higher (because of greater connectivity) or lower (because mainland assemblages are less likely be undersaturated).

Alien species are much more common on human-dominated sites than on more natural sites (especially minimally-used primary vegetation), as expected from their tolerance to disturbed conditions, their colonisation ability and the fact that they may need to displace native species in undisturbed sites [15]. The high abundance of aliens in agricultural sites and cities (Fig 1) has been highlighted previously [25,86–90], and often follows from intentional introduction (e.g., for livestock forage [91]; or, in cities, for trade [92] and ornamental gardening [93]). Overall, land-use change boosted alien abundance more than alien richness, suggesting that those alien species able to tolerate the extreme conditions of disturbed sites can become very successful and abundant [94,95] even if few species are introduced and become established – perhaps also partly because island natives are poor competitors in human-dominated land uses [15].

Sites on more isolated islands have higher absolute numbers of alien species across most land uses (Fig 3) and higher alien abundance in primary vegetation (Fig 10 in S1 File). These results support recent evidence that island remoteness promotes invasions by alien species worldwide [42], and provide further evidence that island biogeography of exotic species is not defined by the natural species–isolation relationship [40,41]. Redding et al. [31] recently found that the presence of other groups of introduced species at the introduction location is one of the main determinants of successful establishment, suggesting an important role for location-specific factors. The reduced diversity of remote islands [39] may make them more invasible [41,96] (but see [97]) because of their high resource availability, missing functional groups and/or low pressures from competitors, pathogens or predators [16,98]. Isolated islands are expected to have particularly small species pools because of restricted immigration [39], leading to lower species richness per unit area and smaller samples from the set of potential species that can survive in different conditions [18]. Through its influence on the size of the species pool, isolation may tip the balance between natives and aliens in terms of which species colonise disturbed sites. We found that alien richness and abundance in minimally-disturbed primary vegetation decreases rapidly as surrounding landmass (and hence the source pool of natural colonists) increases, suggesting that saturated native assemblages are able to resist the influx of aliens better [99].

Factors related to the arrival or introduction of alien species are also likely to partly drive the high alien richness in human-dominated sites on isolated islands. In particular, propagule and/or colonisation pressure – positively related to the establishment of aliens [11,31] – might be higher in remote islands [40]: because such islands often have few native species that can be used for farming, hunting, as sources of fuel or for other economic purposes [16,40], there may be more intentional releases of alien species and high levels of imports [17].

Conversion of minimally-used primary vegetation to other land uses affects composition of alien assemblages more dramatically than native assemblages (Fig 4; Fig 15 in S1 File): the low number and abundance of alien species in natural habitats (Figs 7 and 8 in S1 File) means that other land uses are less likely to have alien species in common with minimally-disturbed sites. The native assemblages at disturbed sites on islands are likely to be nested subsets of undisturbed communities, given the small source pool of potential native colonists, their lack of adaptation to disturbed sites, and their poor competitiveness [16,18,100]. Pastures seem to be an important exception, including some native novel species (i.e., native species that are not present in minimally-disturbed vegetation). The pasture sites that are compared with minimally-used primary vegetation are mostly (147 out of 155 sites) not used intensively, and ∼60% of these sites are from the South Island of New Zealand, Australia, Madagascar and Tierra del Fuego, where rangeland (i.e., ecosystems where the native vegetation has the potential to be grazed [101]) is common [102]. Resilient species among the natives may be able to establish in such rangelands and other low-intensity grazed sites.

Most land uses have lower spatial beta diversity than minimally-disturbed sites, for both alien and native assemblages (Fig 5; Fig 16 in S1 File). This biotic homogenization is stronger for alien assemblages only in plantations and, for richness-based estimates (*J*_*R*_), urban sites, in line with previous studies of these particular human-dominated land uses [103,104]. Our results therefore do not suggest that the gain of alien species in disturbed sites is the main cause of assemblage homogenization on islands – a contrast with previous studies mainly focusing on mainlands [104,105]. Instead, homogenization of island assemblages also seems to result from an important reduction of spatial beta diversity for native assemblages across most land uses. Addition of alien species drives homogenization if the same species become widespread across disturbed sites [36]. Our results suggest that such homogenization may not only be occurring in highly disturbed habitats such as pastures and urban sites, but also in forested land uses (i.e., particularly in plantations and primary vegetation with light or intense use): these are likely to harbour few alien species (since competition is higher than in more disturbed habitats) that are shared across sites. In contrast, alien assemblages in croplands are not more homogeneous than assemblages within minimally-disturbed sites, suggesting that the alien species in this land use are not ubiquitous across sites [36,106], perhaps due to different aliens establishing with different crops or different agricultural methods excluding different species depending on their ecology. The homogenization pattern for native assemblages across land uses might be the result of a subtractive homogenization, caused by the loss or decline of different species from different sites [36] and the persistence of common species.

Because we were not able to classify all species as alien or native, our dataset includes a restricted number of islands. Expanding the dataset to more islands would increase the variation in island characteristics, thus increasing the power to detect any effects they have. However, it is also possible that the characteristics used in these models truly do not have an important effect on alien diversity: e.g., any effect of island area on the site-level richness and abundance of alien species might simply be outweighed by anthropogenic factors such as colonisation pressure. High economic activity is expected to increase the probability of arrival and establishment of alien species [10,11,107], which we did not find; but the country-level values of per-capita GDP may not be a good reflection of the economic activity of the islands.

Further limitations of our data include taxonomic and geographic biases, which may bias our inferences of the impacts of land use on island ecological assemblages. Among major taxonomic groups, our data had the highest number of studies for invertebrates, the lowest for plants, and no studies of fungi (Table 5 in S1 File). Studies in Asia, Europe and Oceania were equally common, but our data included fewer studies in Africa and the Americas.

Our models suggest that natives can be replaced by alien species, some of which might be invasive, in degraded habitats [81,108] and that, on islands, species turnover caused by land-use change might be driven mainly by novel alien species (unique to human-dominated land uses) rather than novel native species (e.g., from other ecosystems on the island), since these are unlikely to colonize anthropogenic habitats. Although this remains speculative, it is likely that mainland settings, having larger species pools than islands, could provide a larger number resilient natives able to establish and compete in disturbed sites [18]. Similar studies of mainlands would be able to assess whether the effects of land-use change on mainland assemblage composition may be driven by both alien and native synanthropic species missing from natural habitats but assembling into novel human-dominated ecosystems.

A question that remains is whether land-use change is the main driver of native species decline or if its effects interact with the presence of alien species [109]. We did not account for the effects of alien richness or abundance on native communities, but previous studies have highlighted habitat modification as the main driver of biodiversity decline, outcompeting other drivers such as invasive species or climate change [83,110]. However, on islands, alien species are important drivers of change and native extinctions [111,112]; therefore, more comprehensive analyses evaluating interactions between different human pressures are needed to disentangle the importance of different threats in driving losses of island biodiversity.

## Acknowledgements

We thank Jason Tylianakis for his advice on compositional similarity models and Tim Newbold, Luca Börger and Jörn Scharlemann for their statistical advice. We are grateful to UNEP-WCMC for sharing data from the Global Island Database and to Samantha Hill for helping with its use. We also thank Dena Spatz for helping with the assemblage of data from the Threatened Island Biodiversity database, and Christian König for contributing to the revision of the manuscript. We are grateful to all data contributors to the PREDICTS database and all contributors for the classification of species as aliens or natives. We thank all current and past members of the PREDICTS projects for their work in data collation and curation.

## Supporting information

**S1 File. Supporting figures and tables**.

