## Supplementary material for "Effects of land-use change and related pressures on alien and native subsets of island communities": S1 File. Supporting figures and tables

### S1 File. Supporting figures and tables for “Effects of land-use change and related pressures on alien and native subsets of island communities”

Table 1. External sources that were used to determine the status of species in the PREDICTS database at either island or country level.

| Source acronym | Citation/Description | Access to data | Species status in |
|---|---|---|---|
| AGDAWR | Australian Government Department of Agriculture and Water Resources. | Search for specific species at: http://www.agriculture.gov.au/ [Accessed March 2017] | Country |
| ALA | Atlas of Living Australia. | Search for specific species at: https://www.ala.org.au/ [Accessed March 2017] | Country |
| AntMaps | Janicki J, Narula N, Ziegler M, Guénard B, Economo EP. Visualizing and interacting with large-volume biodiversity data using client–server web-mapping applications: The design and implementation of antmaps.org. Ecol Inform. 2016;32:185–93. | Search for specific species at: http://antmaps.org/ [Accessed March 2017] | Country |
| AntWeb | AntWeb, Version 8.0.5. Database: AntWeb [Internet]. Available https://www.antweb.org | Search for specific species at: https://www.antweb.org [Accessed May 2017] | Country |
| BirdLife | BirdLife Australia. | Downloaded from: http://birdlife.org.au/conservation/science/taxonomy [Accessed April 2015] | Country |
| | BirdLife International. | Downloaded from: http://datazone.birdlife.org/species/search [Accessed March 2017] (Search for endemic species in countries included in PREDICTS database) | Country |
| C.Vink, Canterbury Museum, NZ | Taxonomic expert at Canterbury Museum, New Zealand | Direct contact with taxonomic expert to classify species from the order Araneae in New Zealand | Country |
| CITES | UNEP. The Species + Website. Nairobi, Kenya. Compiled by UNEP-WCMC, Cambridge, UK. Database: Species + [Internet]. Available from: https://www.speciesplus.net/species | Downloaded from: https://www.speciesplus.net/species [Accessed March 2017] | Country |
| CMS | | | |
| DLO | Discover Life. | Search for specific species at: http://discoverlife.org [Accessed March 2017] | Country |
| ORIG-NATIVE | Data from Flora Europaea (http://rbg-web2.rbge.org.uk/FE/fe.html) and different web APIs. | Data accessed through R package 'originr' function 'is_native') [Accessed March 2019] | Country |
| GAVIA | Dyer EE, Redding DW, Blackburn TM. The global avian invasions atlas, a database of alien bird distributions worldwide. Sci Data. 2017;4(1):170041. | Downloaded from: https://figshare.com/articles/Data_from_The_Global_Avian_Invasions_Atl | Country |

| | | | |
|---|---|---|---|
| | | as_-_A_database_of_alien_bird_distributions_worldwide/4234850 [Accessed May 2017] | |
| GIFT | Weigelt P, König C, Kreft H. GIFT – A Global Inventory of Floras and Traits for macroecology and biogeography. J Biogeogr. 2020;47:16–43. | Data provided by the GIFT team (July, 2017) | Island |
| GISD | ISSG. The Global Invasive Species Database, Version 2015.1. Database: Global Invasive Species Database [Internet]. Available from: http://www.iucngisd.org/gisd/ | Data accessed through R package 'originr' function 'gisd' [Accessed March 2019] | Country |
| GloNAF | van Kleunen M, Pyšek P, Dawson W, Essl F, Kreft H, Pergl J, et al. The Global Naturalized Alien Flora (GloNAF) database. Ecology. 2019;100(1):e02542. | Data provided by the GloNAF team (July 2017) | Island |
| GloNAF-country | | Data provided by the GloNAF team (September 2015) | Country |
| GRIIS | Pagad S, Genovesi P, Carnevali L, Schigel D, McGeoch MA. Introducing the Global Register of Introduced and Invasive Species. Sci Data. 2018;5(1):170202. | Downloaded from: http://www.griis.org/ [Accessed May 2017] (Selected terms: "terrestrial", "freshwater", "verified record") | Country |
| IUCN | IUCN. IUCN Red List of Threatened Species, Version 2017-1. Database: The IUCN Red List of Threatened Species [Internet]. Available from: https://www.iucnredlist.org | Data accessed through R package 'rredlist' function 'rl_occ_country' [Accessed March 2019] | Country |
| Knight (1974) | Knight WJ. Leaf hoppers of New Zealand: Subfamilies Aphrodinae, Jassinae, Xestocephalinae, Idiocerinae, and Macropsinae (Homoptera: Cicadellidae). New Zeal J Zool. 1974;1(4):475–493. | Data taken from publication | Country |
| LCR-NZ | Landcare Research New Zealand. | Search for specific species at: https://www.landcareresearch.co.nz/science/plants-animals-fungi [Accessed May 2017] | Country |
| N.Wyatt, NHM, UK | Taxonomic expert at the Natural History Museum, United Kingdom. | Direct contact with taxonomic expert to classify species from the order Diptera in different countries | Country |
| NZTCS | New Zealand Threat Classification System. Database: NZ Threat Classification System (NZTCS) [Internet]. Available from: https://nztcs.org.nz | Search for specific species at: https://nztcs.org.nz/ [Accessed March 2017] | Country |
| OSF | Orthoptera Species File, Version 5.0. Database: Orthoptera Species File Online [Internet]. Available from: http://orthoptera.speciesfile.org/HomePage/Orthoptera/HomePage.aspx | Search for specific species at: http://orthoptera.speciesfile.org/HomePage/Orthoptera/HomePage.aspx [Accessed May 2017] | Country |
| TEARA | Te Ara: The encyclopedia of New Zealand. | Search for specific species at: https://teara.govt.nz/en [Accessed March 2017] | Country |
| TIBD-IAS | TIB Partners. The Threatened Island Biodiversity Database: developed by Island Conservation, University of California Santa | Data provided by the TIB team (May 2017) (Invasive species on islands) | Island |

| | | | |
|---|---|---|---|
| TIBD-TSpI | Cruz Coastal Conservation Action Lab, BirdLife International and IUCN Invasive Species Specialist Group, Version 2017. Database: The Threatened Island Biodiversity Database [Internet]. Available from: http://tib.islandconservation.org | Data provided by the TIB team (May 2017) (Threatened species on islands) | Island |
| WCSP | World Checklist of Selected Plant Families. | Search for specific species at: http://wcsp.science.kew.org [Accessed March 2017] | Country |
| WPB | Checklist of the Western Palaearctic Bees (Hymenoptera: Apoidea: Anthophila). | Search for specific species at: http://westpalbees.myspecies.info. [Accessed March 2017] | Country |

Table 2. Number of biodiversity records per species status for island data in the PREDICTS database.

| Status | Number of records |
|---|---|
| Alien | 89472 |
| Native | 607586 |
| Not classified | 642281 |

Table 3. Number of biodiversity records that were classified by external sources or data sources in the PREDICTS database. The table shows the number of records classified by each external source (first row shows the total records classified by all external sources), but only the total records classified by all PREDICTS data sources (bottom row). Full names of the external sources are provided in Table 1. If a source in Table 1 is not listed, there were no matches for the species-country combinations in that data source or matching combinations were classified first by other sources.

| Source | Records |
|---|---|
| All external sources | 281981 |
| AGDAWR | 52 |
| ALA | 20 |
| AntMaps | 281 |
| AntWeb | 1934 |
| BirdLife | 33704 |
| C.Vink, Canterbury Museum, NZ | 1830 |
| CITES | 10399 |
| CMS | 735 |
| DLO/NZH | 86 |
| GAVIA | 3833 |
| GIFT | 122833 |
| GISD | 13612 |
| GloNAF | 11508 |
| GloNAF-Nature | 1165 |
| GRIIS | 16497 |
| IUCN | 58467 |
| Knight (1974) | 20 |
| LCR-NZ | 360 |
| N.Wyatt, NHM, UK | 2326 |
| NZTCS | 236 |
| OSF | 754 |
| TEARA | 360 |
| TIBD-IAS | 440 |
| TIBD-TSpl | 379 |
| WCSP | 10 |
| WPB | 140 |
| PREDICTS data sources | 415077 |

Table 4. Variability of different attributes of PREDICTS studies that were included in our analyses (i.e., island data discarding records for species that could not be classified). The first four rows show the minimum/maximum values and the median of calculations performed for each study (e.g., N sites= number of sites sampled in a study). The last two rows show values calculated across all sites of all studies. LMLE= largest maximum linear extent (in meters) across sites within a study (MLE is the length of the maximum distance between multiple sampling points within a site).

| | min | max | median |
|---|---|---|---|
| N sites | 2 | 529 | 18 |
| N species sampled | 1 | 893 | 21 |
| Sampling start | 1986 | 2015 | 2006 |
| LMLE | 0.15 | 14142.136 | 95 |
| Site's species richness | 0 | 135 | 3 |
| Site's total abundance | 0 | 96337 | 7 |

Table 5. Number of sites across UN regions including data for major taxonomic groups (for island data that was used in our analyses). Numbers in brackets show the number of studies (first value) and islands (second value) from which data came from. One island can include several studies and in some cases, a single study sampled several islands.

| Taxon | Africa | Americas | Asia | Europe | Oceania |
|---|---|---|---|---|---|
| Vertebrates | 606 (10; 5) | 63 (3; 3) | 721 (23; 16) | 0 (0; 0) | 853 (21; 18) |
| Invertebrates | 82 (2; 2) | 22 (4; 11) | 166 (12; 7) | 2519 (37; 16) | 935 (22; 4) |
| Plants | 300 (1; 1) | 249 (3; 3) | 400 (10; 5) | 354 (10; 4) | 448 (5; 3) |

Table 6. Number of native and alien species (by major taxonomic group) included in the PREDICTS island data that could be classified. Numbers in brackets show the number of studies (first value) and islands (second value) including data for species of each major taxonomic group.

| Taxon | Native species | Alien species |
|---|---|---|
| Vertebrates | 1248 (57; 42) | 112 (38; 22) |
| Amphibia | 107 | 2 |
| Aves | 794 | 84 |
| Mammalia | 183 | 22 |
| Reptilia | 164 | 4 |
| Invertebrates | 2125 (69; 39) | 384 (33; 17) |
| Annelida | 13 | 1 |
| Arachnida | 63 | 51 |
| Archaeognatha | 3 | 0 |
| Blattodea | 2 | 0 |
| Chilopoda | 12 | 2 |
| Coleoptera | 1315 | 185 |
| Collembola | 1 | 0 |
| Dermaptera | 1 | 2 |
| Diplopoda | 5 | 13 |

| | | |
|---|---|---|
| Diptera | 190 | 26 |
| Hemiptera | 54 | 23 |
| Hymenoptera | 232 | 26 |
| Lepidoptera | 73 | 33 |
| Malacostraca | 5 | 0 |
| Mollusca | 45 | 9 |
| Neuroptera | 3 | 0 |
| Odonata | 61 | 0 |
| Onychophora | 2 | 0 |
| Orthoptera | 30 | 5 |
| Pauropoda | 1 | 0 |
| Psocodea | 8 | 5 |
| Thysanoptera | 4 | 3 |
| Trichoptera | 2 | 0 |
| Plants | 2144 (29; 16) | 298 (27; 14) |
| Bryophyta | 3 | 0 |
| Equisetopsida | 4 | 0 |
| Gnetopsida | 11 | 0 |
| Liliopsida | 549 | 84 |
| Lycopodiopsida | 5 | 0 |
| Magnoliopsida | 1449 | 207 |
| Pinopsida | 18 | 5 |
| Polypodiopsida | 99 | 1 |
| Psilotopsida | 6 | 1 |

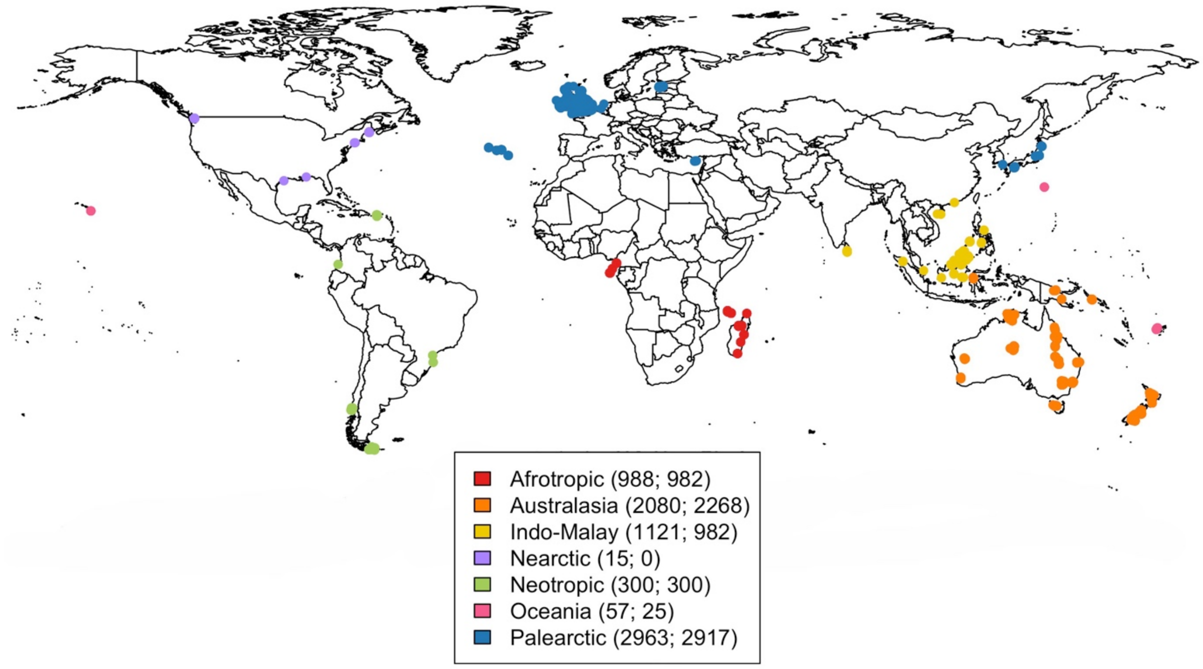


Figure 1. Locations of sites included in the PREDICTS island data that was used for our analyses. Sites are coloured according to the biogeographical realm where they are located. Numbers in brackets show the number of sites per biogeographic realm including data for native (first value) and alien species (second value). Calculations for the number of sites was performed after adding zero measurements for the missing group (aliens or natives) in studies that sampled entire communities (see Methods).

Table 7. Final datasets for the abundance and richness models for alien and native species. The table shows the number of sites, (and in parentheses) studies (first value) and islands (second value) including alien and native data across land use/use intensity categories (LUI). The difference between the number of sites, studies and islands for alien and native data is the result of including data from studies only focusing on sampling a few species (classified as either aliens or natives) either in the native or alien datasets (see Methods).

| | Abundance dataset | | Richness dataset | |
|---|---|---|---|---|
| LUI | Native | Alien | Native | Alien |
| Primary Vegetation Minimal use | 833 (64; 32) | 912 (61; 27) | 1168 (74; 35) | 1247 (71; 30) |
| Primary Vegetation | 534 (34; 26) | 448 (31; 11) | 600 (38; 30) | 514 (35; 15) |
| Secondary Vegetation | 1595 (93; 34) | 1573 (87; 33) | 1824 (103; 37) | 1802 (97; 36) |
| Plantation forest | 675 (40; 21) | 674 (39; 21) | 796 (47; 24) | 795 (46; 24) |
| Cropland | 363 (32; 29) | 330 (29; 14) | 370 (34; 30) | 337 (31; 15) |
| Pasture | 1938 (45; 15) | 2027 (45; 15) | 2135 (48; 17) | 2224 (48; 17) |
| Urban | 491 (14; 6) | 484 (12; 6) | 509 (15; 6) | 502 (13; 6) |

Table 8. Akaike's information criterion (AIC) values for the initial models of total abundance and richness of aliens and natives using the four different random-effects structures that were tested. ∆AIC values are shown relative to the best model. SS= study, SSB= block nested within study, Obs= observation.

| Random-effects structure | d.f. | AIC | ∆AIC |
|---|---|---|---|
| Abundance model | | | |
| (1+LandUse\|SS) + (1\|SSB) + (1\|Island) | 101 | 284.44 | -- |
| (1+LandUse\|SS) + (1\|SSB) | 100 | 294.29 | 9.85 |
| (1\|SS) + (1\|SSB) + (1\|Island) | 74 | 582.15 | 297.71 |
| (1\|SS) + (1\|SSB) | 73 | 591.86 | 307.42 |
| Richness model | | | |
| (1+LandUse\|SS) + (1\|SSB) + (1\|Island) + (1\|Obs) | 101 | 35200.65 | -- |
| (1+LandUse\|SS) + (1\|SSB) + (1\|Obs) | 100 | 35220.26 | 19.61 |
| (1\|SS) + (1\|SSB) + (1\|Island) + (1\|Obs) | 74 | 35404.72 | 204.07 |
| (1\|SS) + (1\|SSB) + (1\|Obs) | 73 | 35427.03 | 226.38 |

**Abundance model**

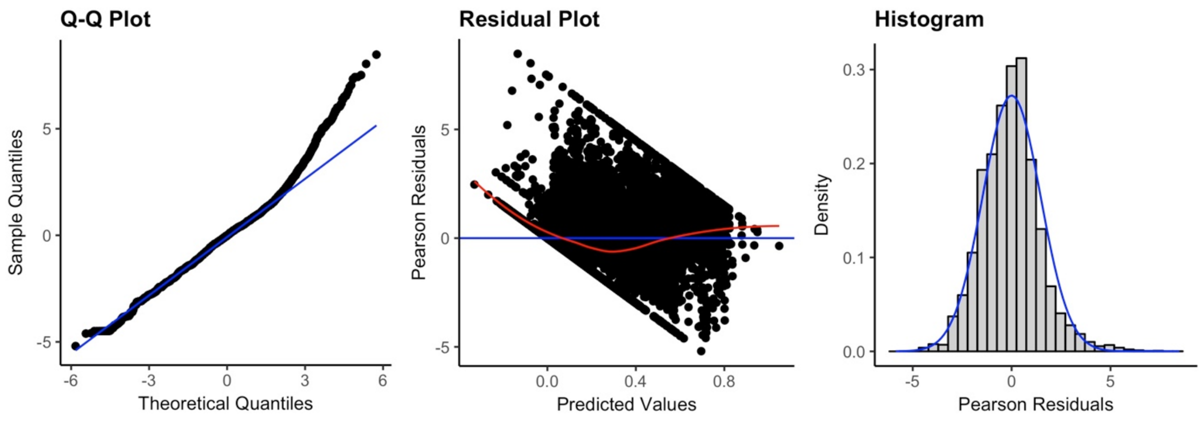


**Species richness model**

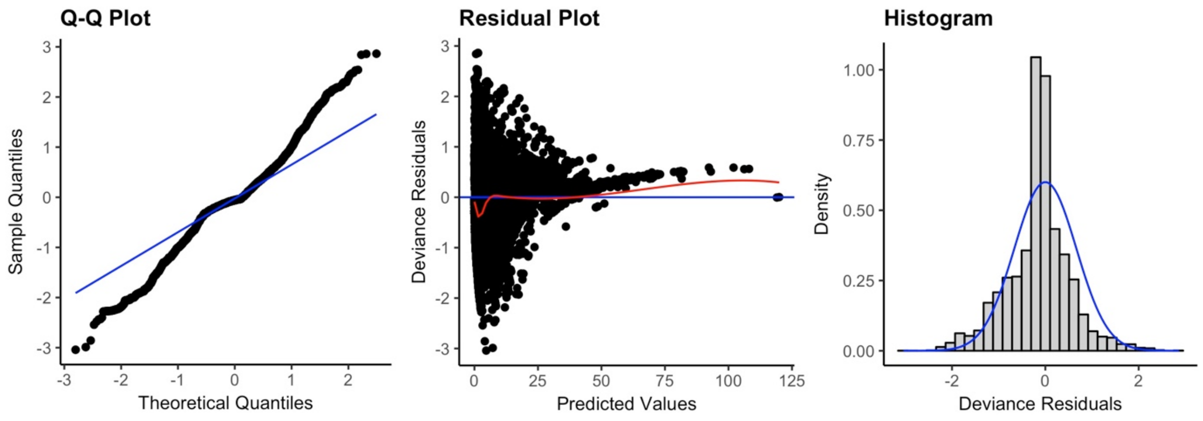

Figure 2. Diagnostic plots for the final models of total abundance and species richness of alien and native species.

Table 9. ANOVA table for the minimum adequate model of total abundance of alien and native species. LUI= land use/use intensity, HPD= human population density, DistRd= distance to the nearest road. Stars indicate the level of significance (Sig): <0.05* and <0.001***

| Term | $\chi^2$ | d.f. | Sig |
|---|---|---|---|
| LUI | 8.641 | 6 | |
| Alien/Native | 10917.632 | 1 | *** |
| HPD | 4.917 | 2 | |
| DistRd | 0.553 | 2 | |
| LUI × Alien/Native | 339.194 | 6 | *** |
| HPD × Alien/Native | 30.713 | 2 | *** |
| LUI × HPD | 25.463 | 12 | * |
| DistRd × Alien/Native | 22.667 | 2 | *** |
| LUI × DistRd | 34.150 | 12 | *** |
| LUI × HPD × Alien/Native | 344.261 | 12 | *** |
| LUI × DistRd × Alien/Native | 86.880 | 12 | *** |

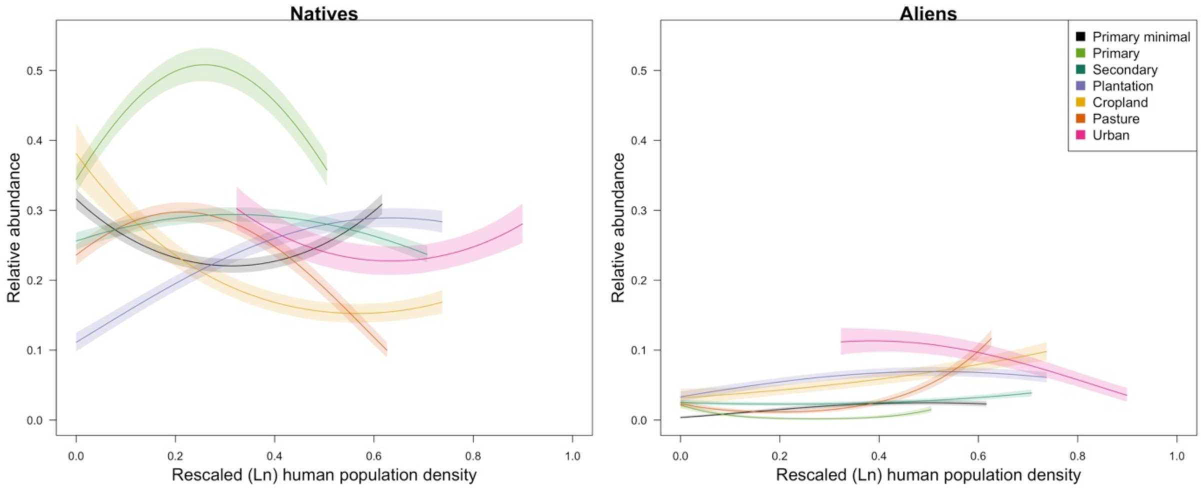


Figure 3 . Response of total abundance of natives and aliens to human population density (HPD) across land uses. The x limits of each coloured line indicate the 2.5th and 97.5th percentiles for the values of HPD represented in each land use in the model dataset. For clarity, the error bars show half the standard error. HPD values are shown on a rescaled axis (as fitted in the models). Abundance is shown on a zero-to-one scale (i.e., abundance rescaled within studies but back-transformed from the square-root scale).

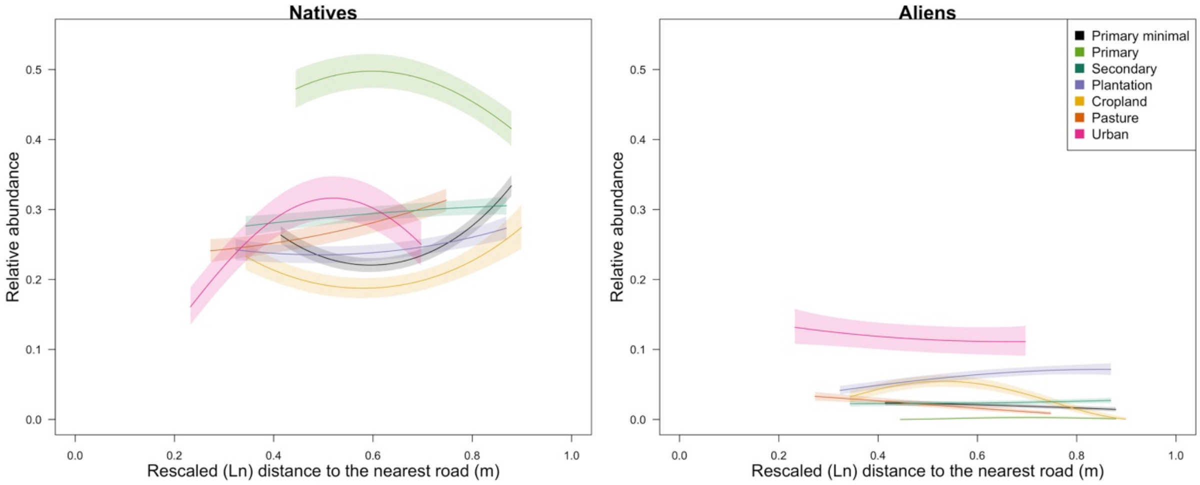


Figure 4. Response of total abundance of aliens and natives to distance to the nearest road (DistRd) across land uses. The x limits of each coloured line indicate the 2.5th and 97.5th percentiles for the values of DistRd represented in each land use in the model dataset. For clarity, the error bars show half the standard error. DistRd values are shown on a rescaled axis (as fitted in the models). Abundance is shown on a zero-to-one scale (i.e., abundance rescaled within studies but back-transformed from the square-root scale).

Table 10. ANOVA table for the minimum adequate model of richness of alien and native species. LUI= land use/use intensity, HPD= human population density, DistRd= distance to the nearest road. Stars indicate the level of significance (Sig): <0.05* and <0.001***

| Term | $\chi^2$ | d.f. | Sig |
|---|---|---|---|
| LUI | 26.292 | 6 | *** |
| Alien/Native | 11588.342 | 1 | *** |
| HPD | 5.061 | 2 | |
| DistRd | 2.484 | 2 | |
| LUI × Alien/Native | 350.059 | 6 | *** |
| HPD × Alien/Native | 299.623 | 2 | *** |
| LUI × HPD | 23.891 | 12 | * |
| DistRd × Alien/Native | 39.828 | 2 | *** |
| LUI × DistRd | 12.982 | 12 | |
| LUI × HPD × Alien/Native | 292.893 | 12 | *** |
| LUI × DistRd × Alien/Native | 76.5157 | 12 | *** |

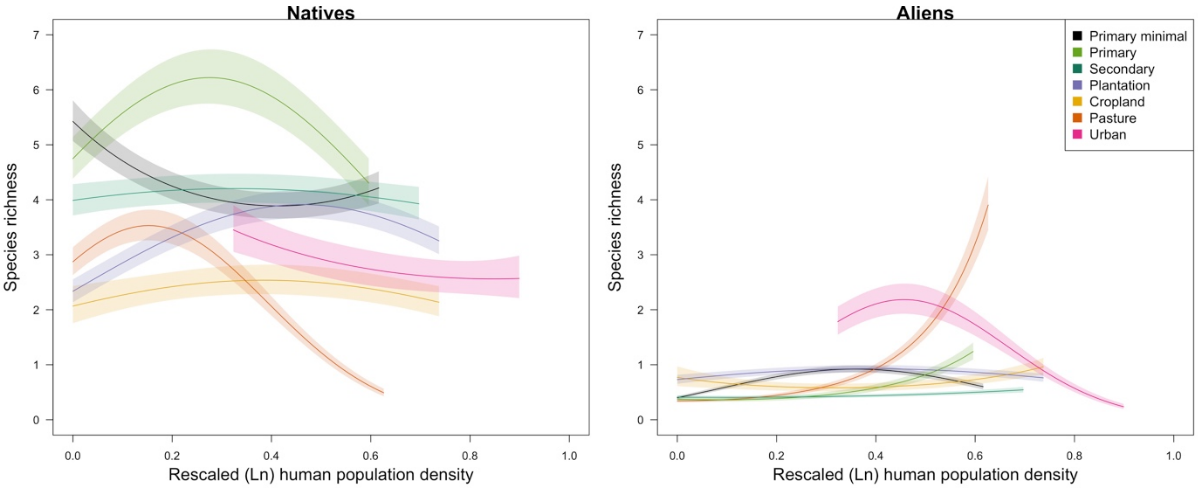


Figure 5. Response of species richness of aliens and natives to human population density across land uses.

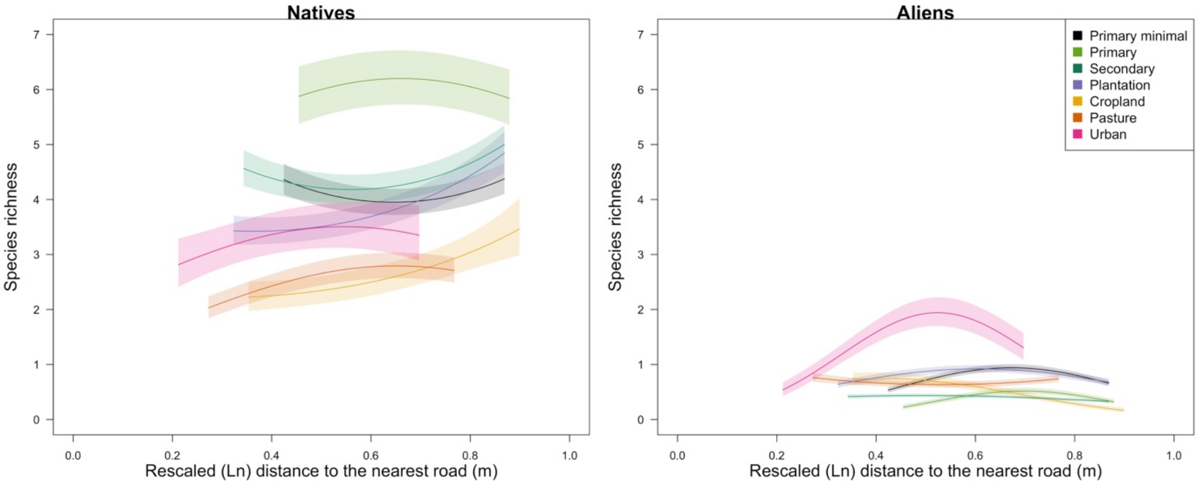


Figure 6. Response of species richness of aliens and natives to distance to the nearest road across land uses.

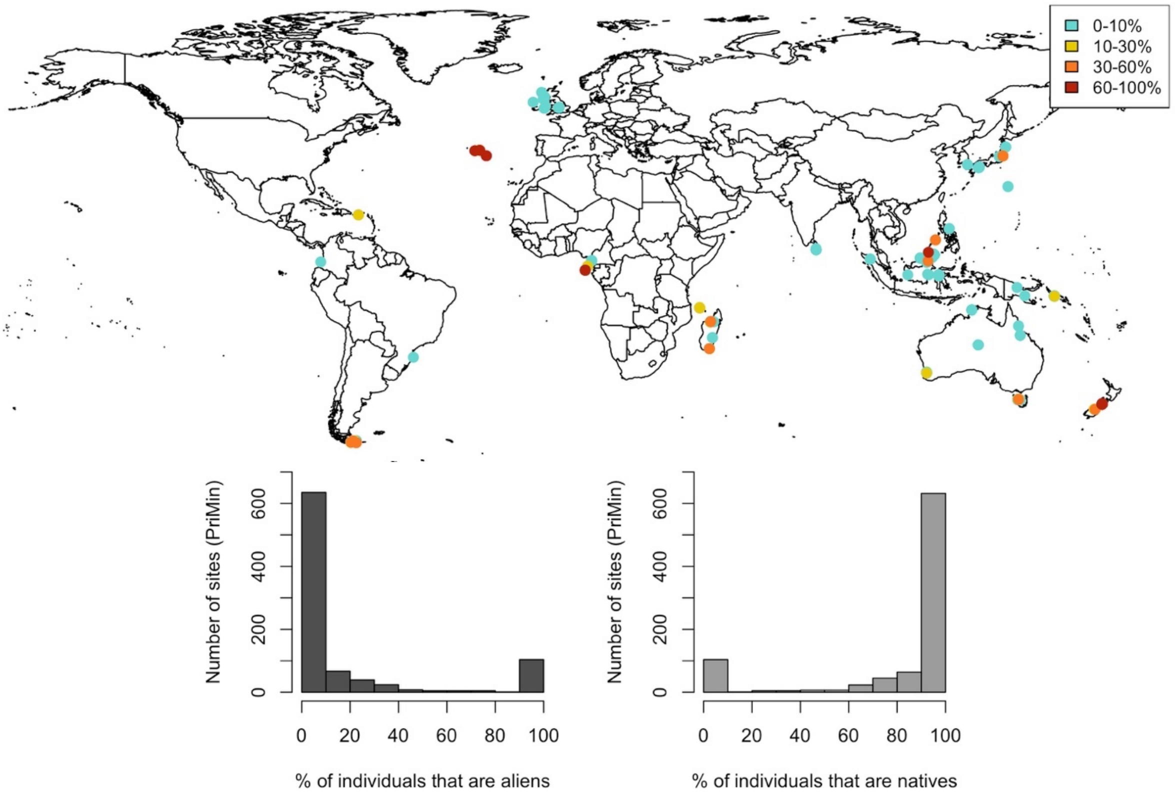


Figure 7. Percentage of individuals that are aliens in sites in minimally-disturbed primary vegetation. Percentages were calculated using exclusively data for alien and native species; species that could not be classified were excluded from these calculations. Only sites that were included in the abundance model are shown. Sites with higher percentages were the last to be plotted, so that they would be highlighted. Histograms show the percentage of individuals that are aliens and natives across sites in minimally-disturbed primary vegetation.

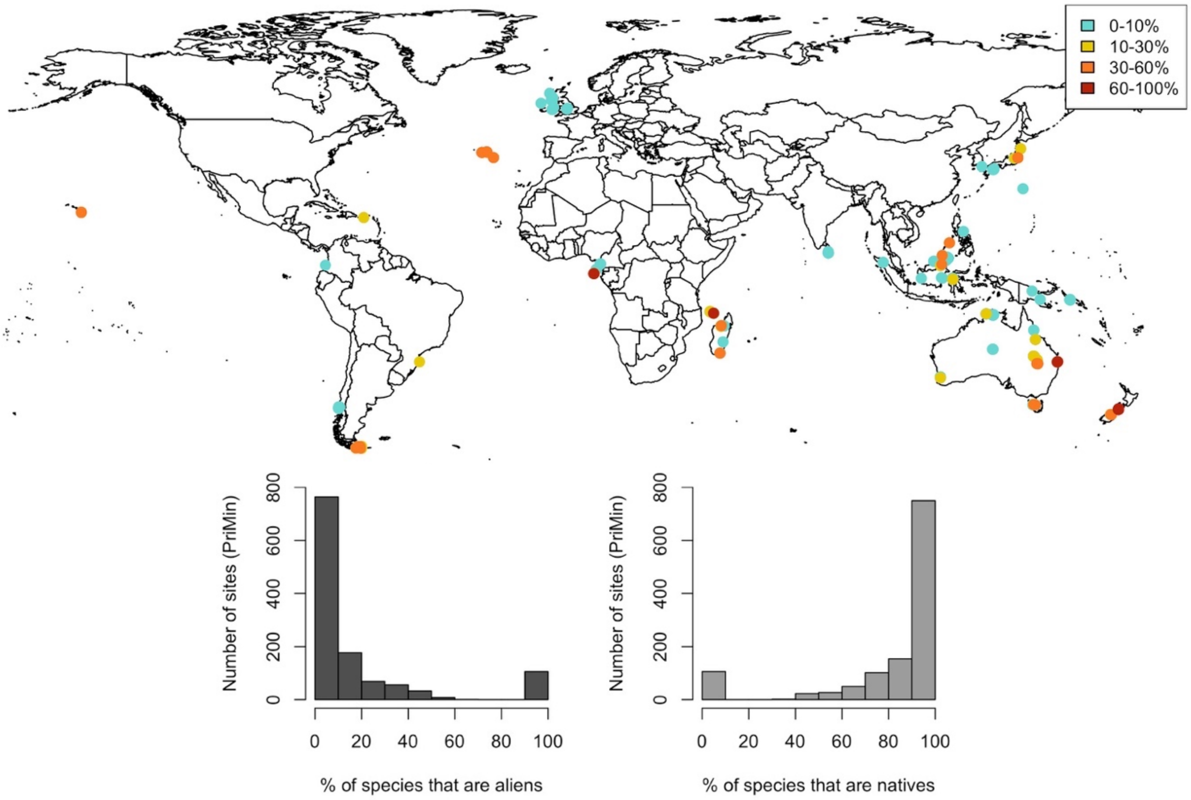


Figure 8. Percentage of species that are aliens in sites in minimally-disturbed primary vegetation. Percentages were calculated using exclusively data for alien and native species; species that could not be classified were excluded from these calculations. Only sites that were included in the richness model are shown. Histograms show the percentage of species that are aliens and natives across sites in minimally-disturbed primary vegetation.

Table 11. Islands included in the models for abundance and richness of aliens using island traits as explanatory variables. Islands marked with a star were included in the abundance and richness models, the rest were only included in richness models. The four islands with missing data for surrounding landmass (i.e., no data in Weigelt et al., 2013) were not included in the models including this variable. In total, the models relating alien abundance to island area or GDP per capita used data from 84 studies in 30 islands, whereas the corresponding species-richness models used data from 98 studies and 39 islands. The model relating landmass to alien abundance used data from 64 studies from 26 islands, whereas the alien richness model used 75 studies from 35 islands. The country listed for each island corresponds to where sites (with data for aliens) in the PREDICTS database are located; i.e., only sites in Borneo were located in two different countries.

| Island | Country | Island area ($km^2$) | Surrounding landmass (summed proportions) | Country per capita GDP (current USD – year 2005) |
|---|---|---|---|---|
| Anijima * | Japan | 7.879 | NA | 37217.649 |
| Anjouan | Comoros | 426.580 | 0.706 | 1068.600 |
| Australia * | Australia | 7588924.738 | NA | 33961.682 |
| Balambangan | Malaysia | 103.175 | 0.704 | 5593.823 |
| Banggi | Malaysia | 431.372 | 0.723 | 5593.823 |
| Bintan | Indonesia | 1169.873 | 0.806 | 1260.929 |
| Borneo * | Malaysia | 732289.104 | 0.501 | 5593.823 |
| Borneo * | Indonesia | 732289.104 | 0.501 | 1260.929 |
| Chichijima * | Japan | 23.757 | NA | 37217.649 |
| Faial * | Portugal | 172.857 | 0.46 | 18784.949 |
| Flores * | Portugal | 140.943 | 0.438 | 18784.949 |
| Grande Comoro * | Comoros | 1015.564 | 0.736 | 1068.600 |
| Great Britain * | United Kingdom | 218670.015 | 0.858 | 41732.641 |
| Hainan * | China | 34023.685 | 0.946 | 1753.418 |
| Hawai'i | United States | 10431.594 | 0.245 | 44307.921 |
| Honshu * | Japan | 227947.264 | 0.573 | 37217.649 |
| Ilha das Rosas * | Brazil | 3.066 | 1.644 | 4770.184 |
| Ireland * | Ireland | 83531.769 | 0.692 | 50878.640 |
| Kolombangara * | Solomon Islands | 693.585 | 0.384 | 880.875 |
| Madagascar * | Madagascar | 587926.700 | 0.46 | 274.820 |
| Mallawalli | Malaysia | 38.321 | 0.785 | 5593.823 |
| New Guinea * | Papua New Guinea | 777319.960 | 0.38 | 770.565 |
| Nishi-jima * | Japan | 0.484 | NA | 37217.649 |
| North Island * | New Zealand | 113707.769 | 0.171 | 27750.725 |
| Osel | Estonia | 2891.685 | 1.548 | 10338.313 |
| Palawan * | Philippines | 11448.371 | 0.494 | 1194.697 |
| Principe * | Sao Tome and Principe | 138.754 | 0.86 | 804.128 |
| Puerto Rico * | Puerto Rico | 8703.443 | 0.381 | 21959.323 |
| Pulau Mangalum | Malaysia | 5.165 | 0.739 | 5593.823 |
| Pulau Mantanai Besar | Malaysia | 2.118 | 0.857 | 5593.823 |

| | | | | |
|---|---|---|---|---|
| Santa Catharina * | Brazil | 422.290 | 1.203 | 4770.184 |
| Santa Maria * | Portugal | 96.926 | 0.459 | 18784.949 |
| Sao Tome * | Sao Tome and Principe | 849.266 | 0.753 | 804.128 |
| South Island * | New Zealand | 150437.674 | 0.163 | 27750.725 |
| Sri Lanka * | Sri Lanka | 65724.996 | 0.569 | 1250.005 |
| Sulawesi* | Indonesia | 168821.235 | 0.51 | 1260.929 |
| Tasmania * | Australia | 63584.062 | 0.346 | 33961.682 |
| Terceira * | Portugal | 400.714 | 0.46 | 18784.949 |
| Tierra del Fuego * | Argentina | 47419.119 | 0.59 | 5076.884 |
| Wight * | United Kingdom | 381.948 | 1.394 | 41732.641 |

Table 12. AIC values for models of total abundance of aliens (including island traits) using the two different random-effects structures that were tested. ΔAIC values are shown relative to the best model. SS= study.

| Random-effects structure | d.f. | AIC | ΔAIC |
|---|---|---|---|
| Model including area | | | |
| (1\|SS) + (1\|Island) | 17 | 555.259 | -- |
| (1\|SS) | 16 | 560.728 | 5.469 |
| Model including surrounding landmass | | | |
| (1\|SS) + (1\|Island) | 17 | -120.244 | -- |
| (1\|SS) | 16 | -117.707 | 2.537 |
| Model including country GDP per capita | | | |
| (1\|SS) + (1\|Island) | 17 | 544.996 | -- |
| (1\|SS) | 16 | 551.138 | 6.142 |

Table 13. ANOVA tables for models of total abundance of aliens including island traits as explanatory variables. LUI= land use/use intensity. Stars indicate the level of significance (Sig): <0.001***

| Term | $\chi^2$ | d.f. | Sig |
|---|---|---|---|
| Model including area | | | |
| LUI | 92.1 | 6 | *** |
| Area | 0.039 | 1 | |
| LUI × Area | 91.771 | 6 | *** |
| Model including surrounding landmass | | | |
| LUI | 95.615 | 6 | *** |
| Landmass | 0.077 | 1 | |
| LUI × Landmass | 97.171 | 6 | *** |
| Model including country GDP per capita | | | |
| LUI | 92.004 | 6 | *** |
| Country GDP | 0.033 | 1 | |
| LUI × Country GDP | 89.801 | 6 | *** |

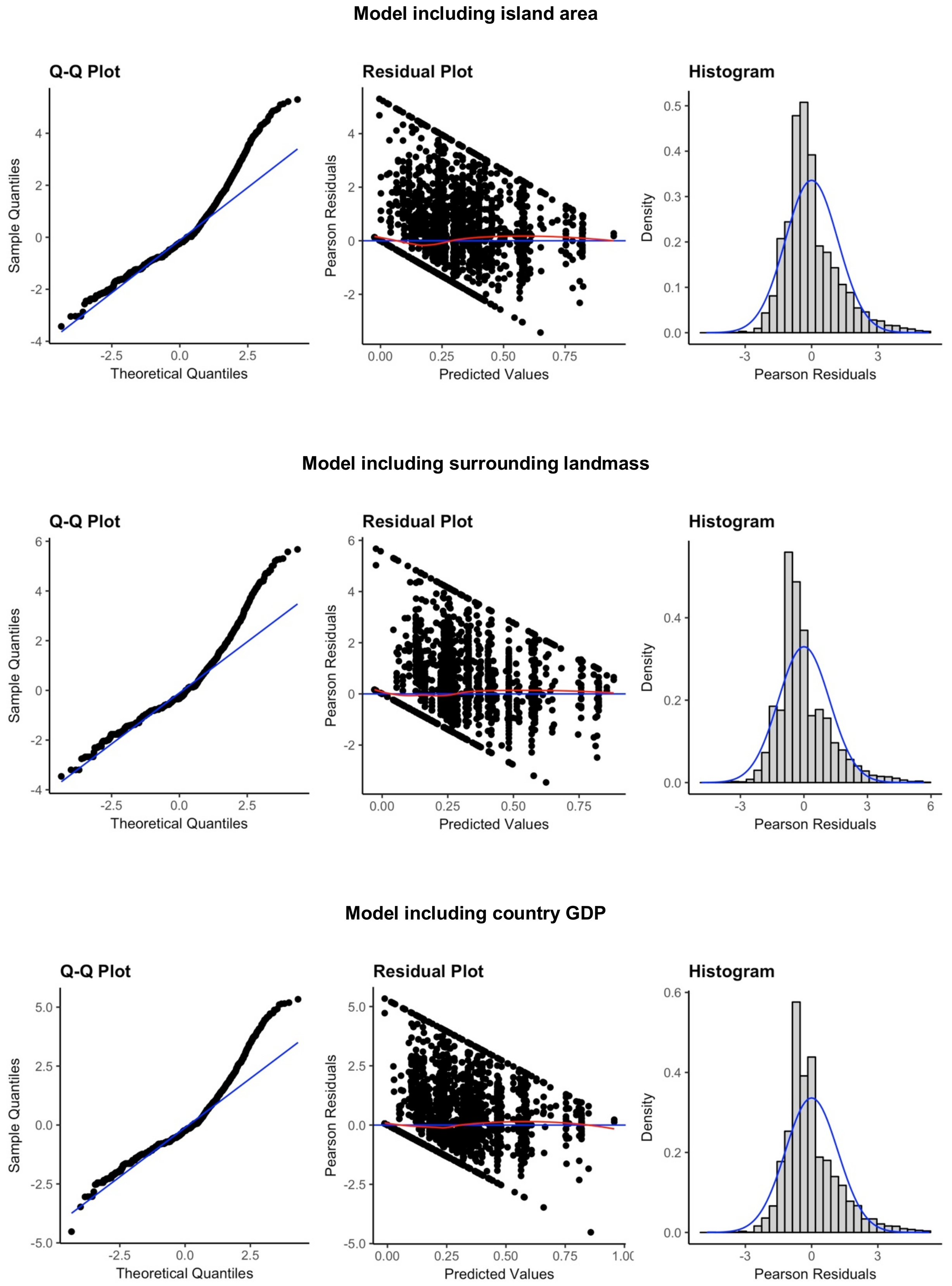


Figure 9. Diagnostic plots for the three models of total abundance of aliens including the different island traits as explanatory variables.

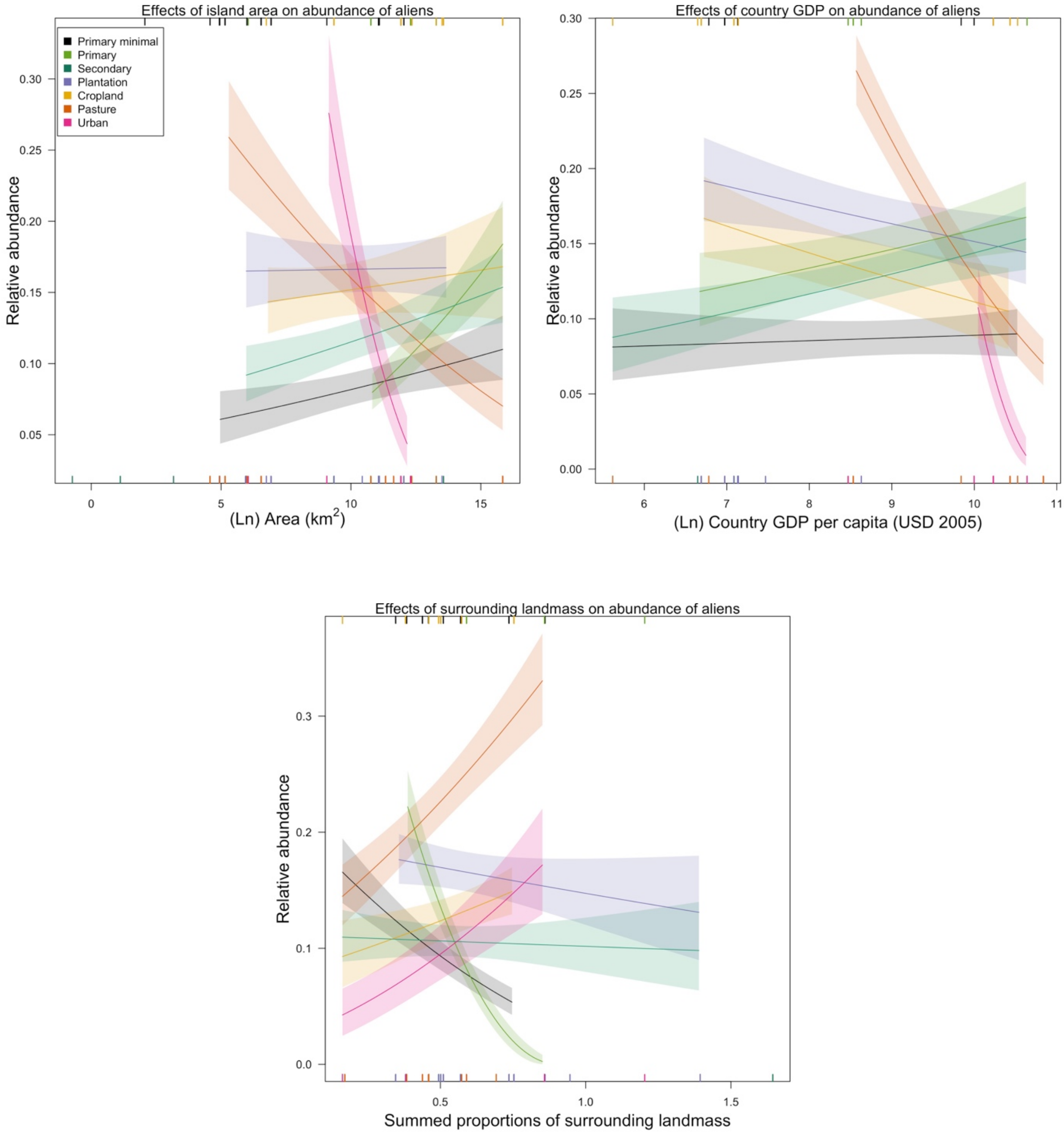


Figure 10. Effects of island area, country-level GDP per capita and surrounding landmass on total abundance of aliens. For clarity, the error bars show half the standard error. Rugs along the horizontal margins show values of the explanatory variables represented (across land uses) in the model data set (rugs for minimally-used primary vegetation, primary vegetation and croplands along the top margin and rugs for the rest of the land uses along the bottom margin). Rugs for land uses can overlap, therefore some data are not visible. The slopes that are significantly different from zero in each model are: Primary (est= 0.03, SE= 0.007, Chisq= 5.5, P= <0.05), Pasture (est= -0.02, SE= 0.005, Chisq= 4.1, P= <0.05) and Urban (est= -0.1, SE= 0.03, Chisq= 8.8 , P= <0.01) in model including island area; Pasture (est= -0.11, SE= 0.01, Chisq= 17.8, P= <0.001) and Urban (est= -0.4, SE= 0.11, Chisq= 12.1, P= <0.001) in model including country GDP; PriMin (est= -0.3, SE= 0.14, Chisq= 4.3, P=<0.05) and Primary Vegetation (est= -0.91, SE= 0.23, Chisq= 13.9, P= <0.001) in model including surrounding landmass

Table 14. AIC values for models of alien species richness (including island traits) using the two different random-effects structures that were tested. ΔAIC values are shown relative to the best model.

| Random-effects structure | d.f. | AIC | ΔAIC |
|---|---|---|---|
| Model including area | | | |
| (1\|SS) + (1\|Island) | 16 | 9300.875 | -- |
| (1\|SS) | 15 | 9344.505 | 43.63 |
| Model including surrounding landmass | | | |
| (1\|SS) + (1\|Island) | 16 | 6555.943 | -- |
| (1\|SS) | 15 | 6624.479 | 68.536 |
| Model including country GDP per capita | | | |
| (1\|SS) + (1\|Island) | 16 | 9315.733 | -- |
| (1\|SS) | 15 | 9397.701 | 81.968 |

Table 15. ANOVA tables for models of alien species richness including island traits as explanatory variables. LUI= land use/use intensity. Stars indicate the level of significance (Sig): <0.01** and <0.001***

| Term | $\chi^2$ | d.f. | Sig |
|---|---|---|---|
| Model including area | | | |
| LUI | 364.134 | 6 | *** |
| Area | 0.006 | 1 | |
| LUI × Area | 74.247 | 6 | *** |
| Model including surrounding landmass | | | |
| LUI | 362.936 | 6 | *** |
| Landmass | 7.136 | 1 | ** |
| LUI × Landmass | 62.746 | 6 | *** |
| Model including country GDP per capita | | | |
| LUI | 360.277 | 6 | *** |
| Country GDP | 0.063 | 1 | |
| LUI × Country GDP | 61.518 | 6 | *** |

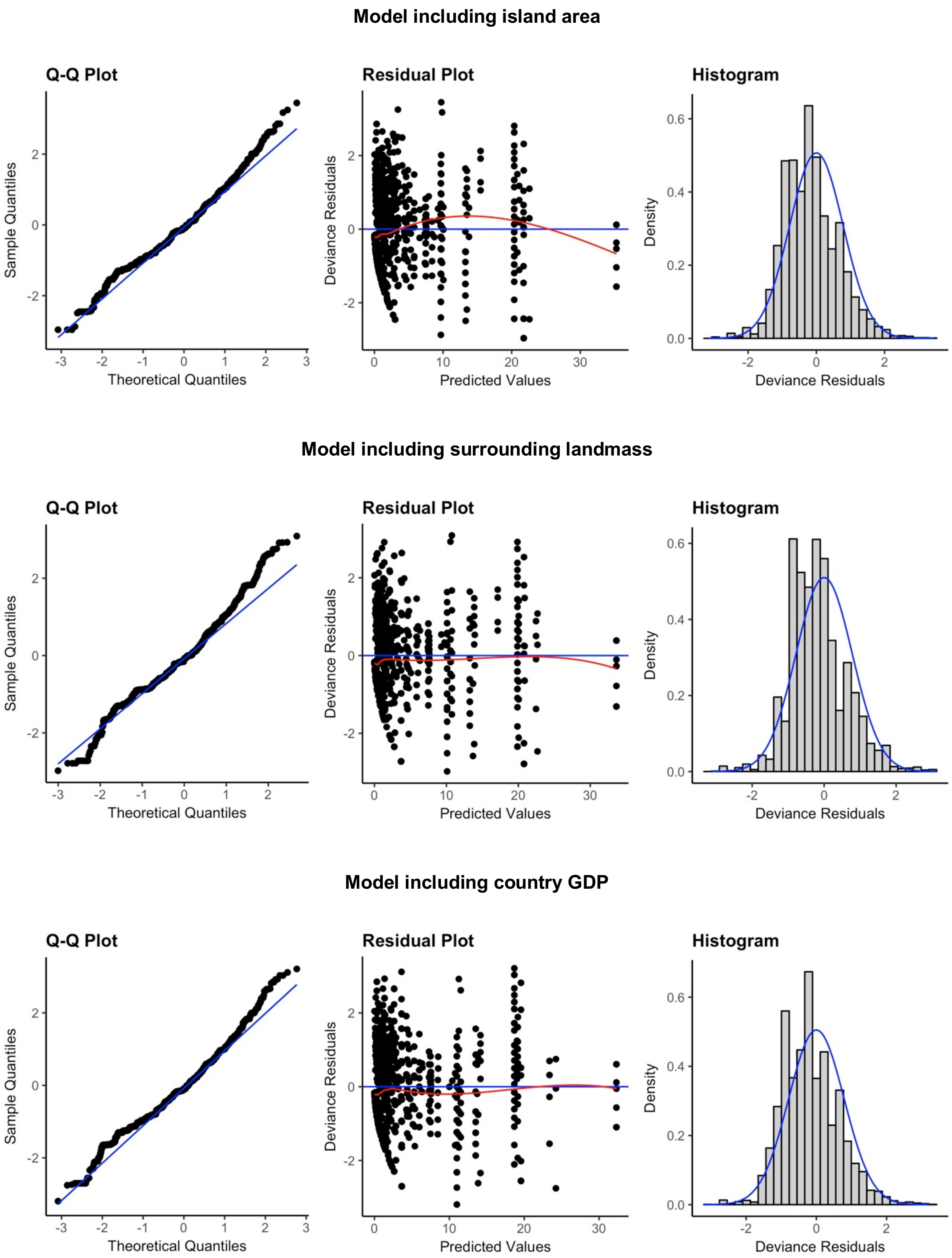


Figure 11. Diagnostic plots for the three models of alien species richness including the different island traits as explanatory variables

.

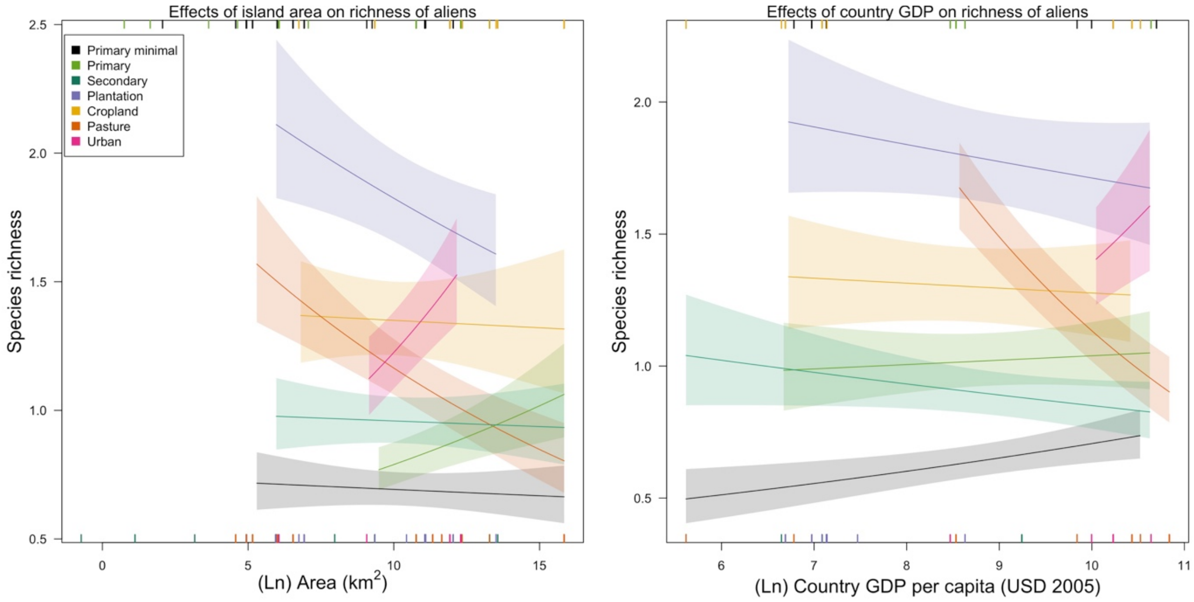


Figure 12. Effects of island area and country-level GDP per capita on species richness of aliens. Rugs along the horizontal margins show values of the explanatory variables represented (across land uses) in the model data set (rugs for minimally-used primary vegetation, primary vegetation and croplands along the top margin and rugs for the rest of the land uses along the bottom margin). Rugs for land uses can overlap, therefore some data are not visible. No slopes were significantly different from zero in the model including island area. In the model including country GDP only Pasture (est= -0.27, SE= 0.05, Chisq= 5.9, P= <0.05) had a slope significantly different from zero.

Table 16. Final dataset for the compositional similarity models for alien and native assemblages. The table shows the number of pair of sites per each land-use contrast generated from pairwise comparisons within studies in the PREDICTS database. Numbers in brackets show the number of studies (first value) and islands (second value) from which data came from. Only land-use contrasts of interest are shown. In total, the models included 91 studies, 53 having data for alien assemblages (from 2,421 sites, 24 islands, and for 591 species) and 89 having data for natives (from 4,274 sites, 32 islands, and for 3,198 species).

| Land-use contrast | Aliens | Natives |
|---|---|---|
| PriMin- PriMin | 10986 (18; 16) | 19053 (35; 20) |
| PriMin- Primary | 4139 (8; 6) | 5627 (12; 7) |
| PriMin- Secondary | 6405 (19; 14) | 12847 (32; 21) |
| PriMin- Plantation | 8994 (11; 8) | 10718 (17; 10) |
| PriMin- Cropland | 2712 (7; 5) | 3477 (9; 6) |
| PriMin- Pasture | 8390 (6; 8) | 9066 (10; 11) |
| PriMin- Urban | 2 (1; 1) | 50 (2; 2) |
| Primary-Primary | 8149 (8; 5) | 22714 (18; 8) |
| Secondary- Secondary | 10326 (26; 15) | 31367 (53; 23) |
| Plantation - Plantation | 28139 (14; 10) | 33374 (22; 14) |
| Cropland - Cropland | 3720 (5; 4) | 4975 (6; 5) |
| Pasture - Pasture | 19006 (14; 11) | 117398 (24; 12) |
| Urban - Urban | 1901 (6; 3) | 14906 (10; 4) |

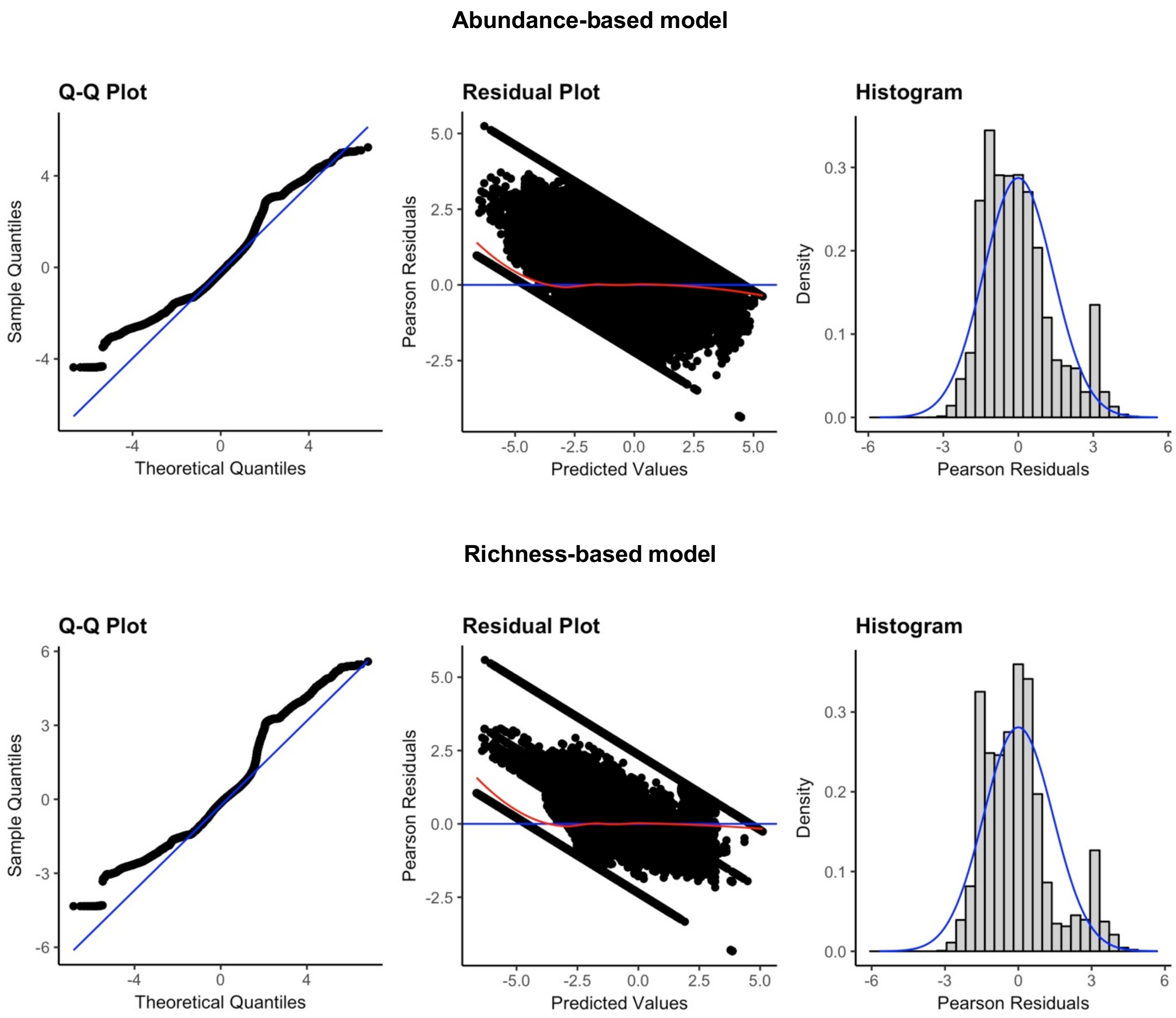


Figure 13. Diagnostic plots for the minimum adequate models of abundance-based and richness-based compositional similarity for alien and native assemblages.

Table 17. Coefficients from the final compositional similarity models (abundance-based and richness-based) for native and alien assemblages. Coefficients for native species (i.e., interaction coefficients) are expressed as the difference from the alien coefficients. Significance (indicated by stars) is shown for the coefficients of interest (first section of the table), for which “two-tailed” tests were performed to compare the observed values against null distributions. Although we only reported significance for the interaction coefficients, we also tested for significance of alien coefficients (baseline in model). The coefficient for the effects of environmental distance on native assemblages is shown as NA for the richness-based model since this variable did not interact significantly with the alien/native term. Significance codes: >0.05 --, <0.05**, and 0.005***

| | Abundance-based model | | Richness-based model | |
|---|---|---|---|---|
| | Aliens | Natives | Aliens | Natives |
| PriMin-PriMin | 0.604 *** | 0.774 *** | 0.073 *** | 0.64 *** |
| Geographic distance | -0.155 *** | 0.019 ** | -0.128 *** | 0.017 ** |
| Environmental distance | -1.371 *** | -0.709 *** | -1.56 *** | NA |
| PriMin-Primary | -0.67 *** | 0.221 *** | -0.486 *** | 0.201 *** |
| PriMin-Secondary | -0.874 *** | 1.175 *** | -0.502 *** | 0.765 *** |
| PriMin-Plantation | -1.82 *** | 2.443 *** | -1.566 *** | 2.093 *** |
| PriMin-Cropland | -2.951 *** | 3.288 *** | -2.743 *** | 3.084 *** |
| PriMin-Pasture | -1.324 *** | 0.382 *** | -0.719 *** | 0.09 ** |
| PriMin-Urban | 0.376 -- | -1.087 -- | -0.134 -- | -0.954 -- |
| Primary-Primary | 0.256 *** | 0.198 *** | 0.495 *** | 0.076 -- |
| Secondary-Secondary | -0.009 -- | 0.369 *** | 0.221 *** | 0.188 *** |
| Plantation-Plantation | 1.428 *** | -0.61 *** | 1.294 *** | -0.528 *** |
| Cropland-Cropland | -0.1 -- | 0.135 -- | -0.047 -- | 0.158 ** |
| Pasture-Pasture | 0.132 *** | 0.347 *** | 0.309 *** | -0.048 -- |
| Urban-Urban | -0.071 -- | 0.118 -- | 0.484 *** | -0.38 *** |
| Cropland-Pasture | 1.331 | -2.909 | 1.674 | -3.138 |
| Cropland-Plantation | -0.558 | 0.396 | -0.345 | 0.168 |
| Cropland-PriMin | -1.195 | 0.343 | -0.96 | 0.108 |
| Cropland-Primary | 2.364 | -2.212 | 2.557 | -2.386 |
| Cropland-Secondary | -0.809 | 0.226 | -0.59 | 0.03 |
| Cropland-Urban | -0.02 | -1.221 | 0.36 | -1.682 |
| Pasture-Cropland | 0.844 | -1.648 | 1.275 | -2.098 |
| Pasture-Plantation | -0.202 | -1.153 | 0.038 | -1.362 |
| Pasture-PriMin | -0.36 | -0.603 | -0.052 | -0.613 |
| Pasture-Primary | -0.083 | -0.455 | 0.117 | -0.409 |
| Pasture-Secondary | -0.624 | -0.444 | 0.065 | -0.894 |
| Pasture-Urban | 0.416 | -0.884 | 0.929 | -1.377 |
| Plantation-Cropland | -1.492 | 2.275 | -1.376 | 2.187 |
| Plantation-Pasture | -0.04 | -0.479 | 0.613 | -1.046 |
| Plantation-PriMin | -0.362 | 0.323 | -0.067 | -0.03 |
| Plantation-Primary | -2.028 | 1.495 | -1.419 | 1.062 |
| Plantation-Secondary | 0.418 | -0.035 | 0.635 | -0.26 |
| Plantation-Urban | 0.321 | -0.693 | 1.087 | -1.688 |
| Primary-Cropland | 0.334 | -0.124 | 0.582 | -0.505 |
| Primary-Pasture | -0.345 | 0.023 | -0.09 | -0.04 |
| Primary-Plantation | -2.258 | 2.446 | -1.877 | 2.107 |
| Primary-PriMin | -0.491 | 0.045 | -0.329 | 0.136 |

| | | | | |
|---|---|---|---|---|
| Primary-Secondary | -0.078 | 0.471 | 0.229 | 0.295 |
| Primary-Urban | 0.364 | -1.655 | 2.607 | -3.695 |
| Secondary-Cropland | -2.53 | 3.321 | -2.355 | 3.183 |
| Secondary-Pasture | -1.197 | 0.293 | -0.351 | -0.125 |
| Secondary-Plantation | -1.113 | 1.903 | -0.832 | 1.547 |
| Secondary-PriMin | 0.246 | -0.228 | 0.406 | -0.466 |
| Secondary-Primary | -0.718 | 0.917 | -0.409 | 0.754 |
| Secondary-Urban | -4.406 | 4.715 | -3.011 | 3.371 |
| Urban-Cropland | 1.118 | -1.197 | 1.783 | -1.787 |
| Urban-Pasture | 0.788 | -1.016 | 1.806 | -1.934 |
| Urban-Plantation | 0.899 | -0.936 | 1.064 | -1.21 |
| Urban-PriMin | 0.842 | -0.988 | 3.112 | -3.241 |
| Urban-Primary | 0.359 | -2.305 | 0.219 | -1.766 |
| Urban-Secondary | -5.243 | 5.048 | -3.723 | 3.592 |

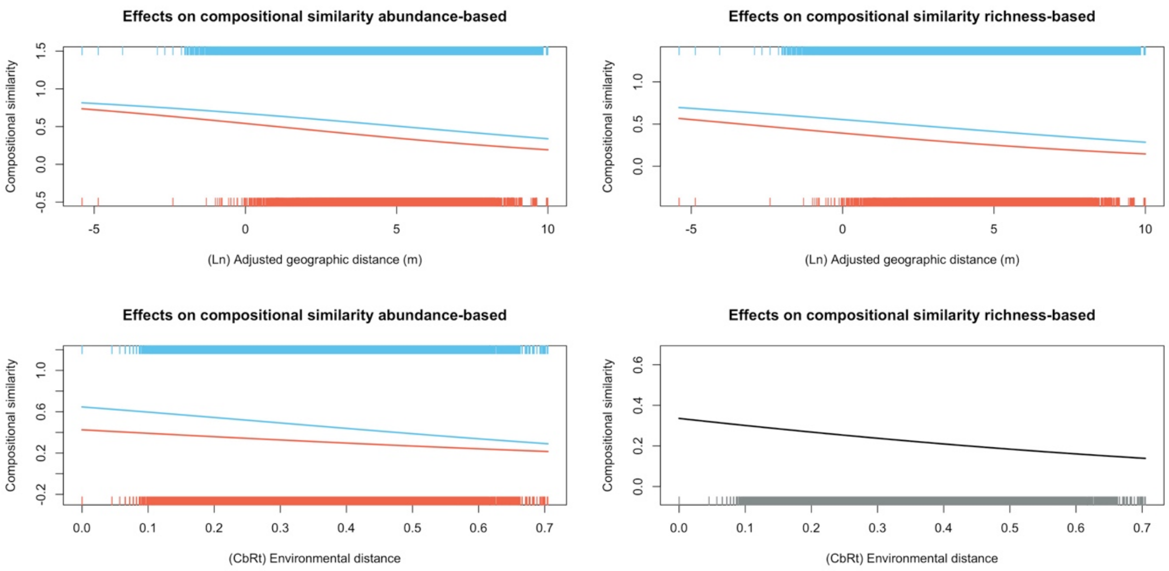


Figure 14. Effects of geographic and environmental distance between sites on compositional similarity ($J_A$ and $J_R$) of alien (orange) and native (blue) assemblages. The rugs in the figures show the distribution of data for aliens and natives. Significance (indicated by stars) corresponds to p-values calculated from "two-tailed" tests using the interaction coefficients (to compare the observed values against null distributions) to test for significant differences between responses of aliens and natives. In the case of the richness-based model, the p-value for environmental distance was calculated for the coefficient of the single term since this variable did not interact significantly with the alien/native term. Significance code: <0.05**, 0.005***

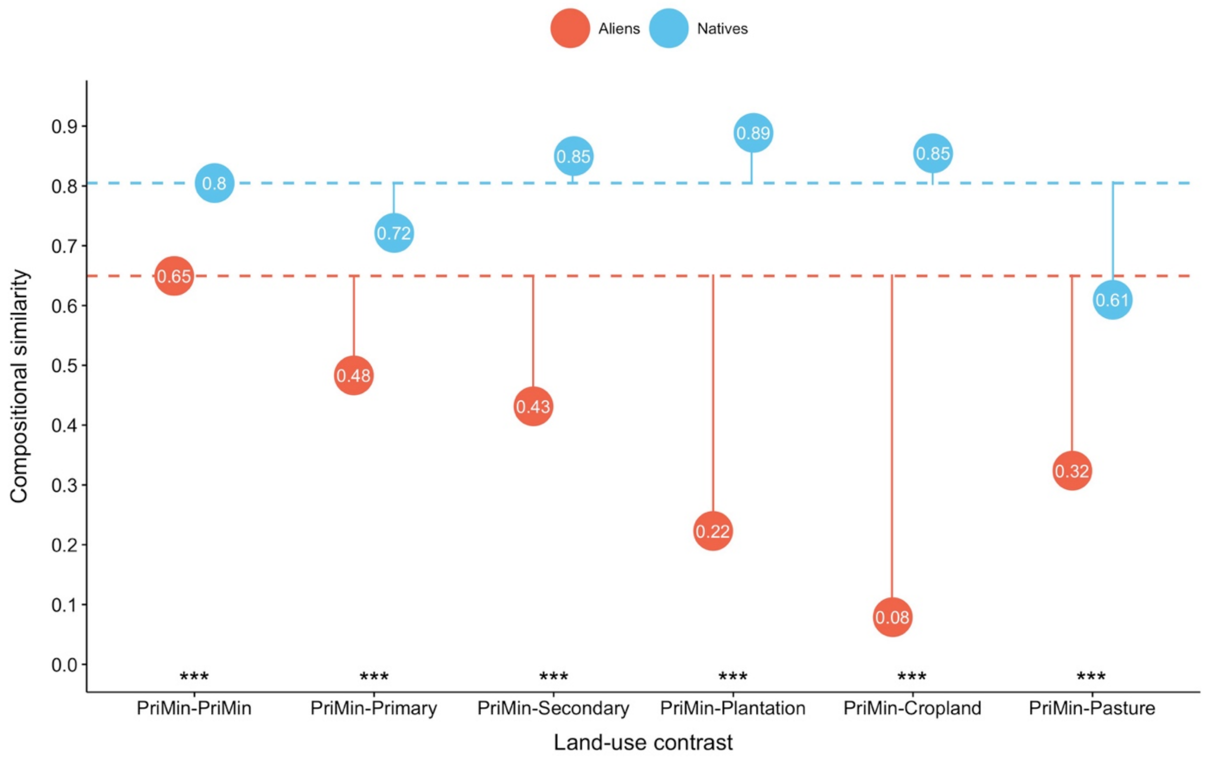


Figure 15. Abundance-based ($J_A$) compositional similarity estimates for land-use contrasts where site *i* is in PriMin. Solid lines show the magnitude of change in $J_A$ driven by change to different land uses; the baseline is compositional similarity between PriMin sites for alien and native assemblages respectively (dashed lines). Significance (indicated by stars) is shown for alien/native differences for $J_A$ changes from PriMin-PriMin on a logit scale (results from "two-tailed" tests comparing the coefficients for interaction between alien/native and land-use contrast to null distributions). Results for the PriMin-Urban contrast are not shown because sample sizes for this contrast were very small (but see the coefficients in Table 16) Significance code: 0.005***

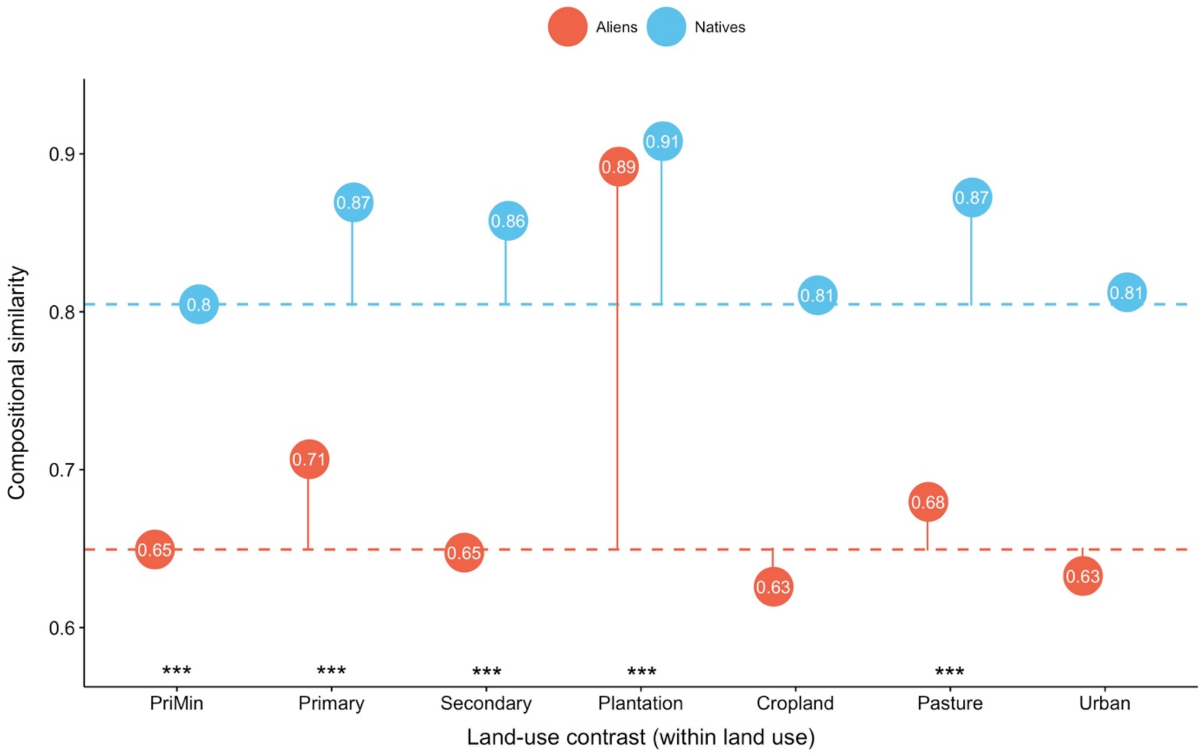


Figure 16. Abundance-based ($J_A$) compositional similarity estimates for alien and native assemblages in sites within the same land use. Each category corresponds to a land-use contrast (i.e., Cropland= Cropland-Cropland). Solid lines show the magnitude of change in $J_A$ using PriMin-PriMin compositional similarity as baseline (dashed lines). Significance connotation and codes as in Figure 15.
